# Expanding the HP1a-binding consensus and molecular grammar for heterochromatin assembly

**DOI:** 10.1101/2024.12.03.626544

**Authors:** Serafin U. Colmenares, Shingo Tsukamoto, Collin Hickmann, Lucy D. Brennan, Mohammad Khavani, Mohammad R. K. Mofrad, Gary H. Karpen

**Author notes:** These authors contributed equally to this work (Serafin U. Colmenares, Shingo Tsukamoto).

## Abstract

The recruitment of Heterochromatin Protein 1 (HP1) partners is essential for heterochromatin assembly and function, yet our knowledge regarding their organization in heterochromatin remains limited. Here we show that interactors engage the Drosophila HP1 (HP1a) dimer through a degenerate and expanded form of the previously identified PxVxL motif, which we now term HP1a Access Codes (HACs). These HACs reside in disordered regions, possess high conservation among Drosophila homologs, and contain alternating hydrophobic residues nested in a cluster of positively charged amino acids. These findings and molecular dynamics simulations identify key electrostatic interactions that modulate HP1a-binding strength and provide a dramatically improved HP1a-binding consensus motif that can reveal protein partners and the molecular grammar involved in heterochromatin assembly. We propose HP1a acts as a scaffold for other heterochromatin components containing HAC motifs, which in turn may regulate the function and higher order structure of the heterochromatin compartment.

## INTRODUCTION

Heterochromatin forms an evolutionarily conserved nuclear compartment critical for genome integrity. In humans and Drosophila, heterochromatin houses 25-30% of the genome, respectively, that is primarily composed of repetitive DNA such as transposons and satellite sequences (Hoskins *et al*., 2007) enriched in gene-poor pericentric and telomeric regions of chromosomes. Important for transposon suppression, chromosome segregation, and telomere protection, heterochromatin displays physical, molecular, and functional attributes distinct from gene-rich euchromatin (Janssen, Colmenares and Karpen, 2018). Constitutively dense through interphase, the heterochromatin compartment is transcriptionally repressive, replicates later in S phase in many organisms, and utilizes alternative DNA repair pathways (Chiolo *et al*., 2011; Janssen, Colmenares and Karpen, 2018).

Various factors converge in heterochromatin to perform its many functions. Transcriptional repression, as assayed by the insertion of reporter genes or chromosome rearrangements, have been attributed to many heterochromatin factors (HP1a, Su(var)3-9, HP2) (Eissenberg *et al*., 1992; Schotta *et al*., 2002; Shaffer *et al*., 2002). Interestingly, hundreds of genes reside in Drosophila heterochromatin, some of which are essential for viability (Weiler and Wakimoto, 1995; Dimitri *et al*., 2003), suggesting the presence of an unknown alternative transcriptional program specific for the heterochromatin landscape. Meanwhile, transcription of repetitive DNA has been shown in many organisms to be instrumental in heterochromatin establishment/maintenance and suppression of transposable elements by providing a non-coding scaffold for small RNAs that recruit heterochromatin complexes (Piwi, Ago2) (Brennecke *et al*., 2007; Czech *et al*., 2008). Heterochromatin also employs factors that inhibit replication fork progression and suppress polytenization (Rif1, SuUR) (Makunin *et al*., 2002; Seller and O’Farrell, 2018), or direct double-strand breaks to the periphery for homologous recombination repair (Smc5/6) (Chiolo *et al*., 2011). Finally, the heterochromatin subnuclear compartment houses components of sister chromatid cohesion (Dmt) (Yamada *et al*., 2017) necessary for the coordinated distribution of sister chromatids during mitosis and undergoes dissolution and re-assembly every cell cycle. How such various factors involved in disparate functions are recruited and organized within the same 3-D heterochromatin compartment remains largely unknown.

Central to heterochromatin organization and function is Heterochromatin Protein 1, or HP1, that specifically binds and propagates the heterochromatic histone mark di-and tri-methylated H3 K9 by recruiting methyltransferases. HP1 also binds multiple proteins, many of which have been shown to require HP1 for enrichment in heterochromatin. HP1 is comprised of two globular structured domains: a chromodomain (CD) that binds methylated H3 K9 (Bannister *et al*., 2001; Lachner *et al*., 2001) and a chromoshadow domain (CSD) important for dimerization and protein partner binding (Brasher *et al*., 2000). Other than short, disordered regions at the flanks of the protein, HP1 also possesses a single unstructured hinge region between the two globular domains that binds nucleic acid (Muchardt *et al*., 2002). Previously, we and others have shown that the Drosophila homolog HP1a co-purifies with dozens to hundreds of proteins (Alekseyenko *et al*., 2014; Swenson *et al*., 2016), and that some partners display a remarkable variation of enrichment patterns, or “subdomains,” that indicate the presence of a higher-order structure within heterochromatin (Swenson *et al*., 2016). Whether the large number of partners utilize individual mechanisms to target HP1a or a common motif to access the heterochromatin domain is an important fundamental question for understanding heterochromatin structure and organization.

A pentameric HP1-binding motif (PxVxL) was previously identified through a phage display screen to bind the HP1 CSD dimer. Based on the peptide consensus sequence, this pentamer contains alternating hydrophobic amino acids of proline, valine, and leucine that give rise to its name (Smothers and Henikoff, 2000). Structural studies showed that PxVxL peptides bind asymmetrically to a hydrophobic pocket formed by the HP1 CSD dimer, with the central valine occupying the cleft formed by the two CSD C-termini (Thiru *et al*., 2004). Other domains of HP1, such as the CD and hinge regions, have also been implicated in binding other proteins (Badugu *et al*., 2005). However, most examples of HP1 partners identified across many species from fission yeast to humans utilize the PxVxL motif, signifying its evolutionary importance (Thiru *et al*., 2004; Coustham *et al*., 2006; Honda and Selker, 2008).

The recent discovery that heterochromatin domains display biophysical properties of biocondensates *in vivo* (Strom *et al*., 2017) has also prompted a re-evaluation of the PxVxL motif as a potentially important contributor to emergent properties and functions. As biocondensates form membraneless compartments through multivalency and the cumulative weak interactions among components, we propose that HP1 acts as a scaffold that recruits multiple heterochromatin components through the PxVxL motif. In addition, peptides containing PxVxL motifs have been shown to enhance HP1 dimerization (Mendez *et al*., 2011) and promote or inhibit phase separation *in vitro* and *in silico* (Thiru *et al*., 2004; Larson *et al*., 2017; Her *et al*., 2022). Thus, the PxVxL motif may represent not only a common HP1a interaction interface but also a mechanism to modulate local or global HP1a dynamics and condensate composition and biophysical properties.

Variants of the PxVxL motif, such as LxVxL, LxVxI, and CxVxL, have also been identified (Bassett *et al*., 2008; Mendez *et al*., 2011; Mendez, Mandt and Elgin, 2013; Meyer-Nava *et al*., 2020). Interestingly, a non-canonical motif (LxVxI) displayed a higher binding affinity to the HP1a CSD dimer than canonical (PxVxL) and near-canonical (PxVxV) motifs (Mendez *et al*., 2011). The canonical PxVxL motif is also not universally sufficient for binding, as evidenced by mammalian STAM2a, which has a PxVxL motif but fails to bind the CSD (Lechner *et al*., 2005). Furthermore, the HP1a C-terminal extension (CTE) and the flanking residues of the PxVxL motif are implicated in HP1a-partner binding selectivity and interaction strength (Thiru *et al*., 2004; Mendez *et al*., 2011; Mendez, Mandt and Elgin, 2013; Liu *et al*., 2017). Deletion of the last 3 residues on the CTE enhanced HP1a dimerization but severely reduced the binding of PxVxL peptides, whereas chemical shift perturbations were detected at HP1a residues that contact positions beyond the PxVxL pentamer core (Mendez *et al*., 2011). Altogether, these findings raise questions about the combination of hydrophobic residues sufficient for PxVxL function, and whether additional molecular features exist beyond the pentapeptide motif.

Here we utilize a combination of evolutionary comparisons, *in vivo* imaging, and full atomistic molecular dynamics (MD) simulations to comprehensively define the molecular signature of the PxVxL motif in Drosophila and determine the specific interactions that regulate HP1a binding. Our analyses show that PxVxL motifs of HP1a-binding partners are highly conserved across Drosophila species, embed in mainly disordered domains of HP1a partner proteins, and are driven by charged residues that we now propose to be a critical component of the molecular grammar for heterochromatin targeting. As a result, we have expanded the number of known functional HP1a-binding sites and provide an improved signature with higher predictive power, which we refer to as the HP1a Access Code, or “HAC”. We have also determined that HACs exhibit a range of affinity for HP1a, which are modulated by electrostatic interactions. We propose that HACs embedded in disordered domains are an important part of the molecular language used to assemble the heterochromatin condensate, and that differences in HP1a affinity among protein partners are critical for the composition, ultrastructure, and function of heterochromatin.

## RESULTS

### Imaging peptide localization in cells identifies new HP1a-binding sequences

Discovery of the foundational HP1a-binding pentamer sequence, PxVxL, showed that it consists of alternating nonpolar residues with the central valine, designated as position 0, binding the junction of two CSD domains in an HP1a dimer (**Figure 1A**). Correspondingly, the proline (P) was designated as position-2 and leucine (L) as position +2, while positions-1 and +1 displayed higher variability, and were thus represented as ‘x’. Together, the trio of nonpolar residues occupy a hydrophobic pocket formed by the HP1a CSD dimer, as shown by structural studies (Thiru *et al*., 2004), and is thought to mediate the ability of HP1a to associate with protein complexes.

**Figure 1.**
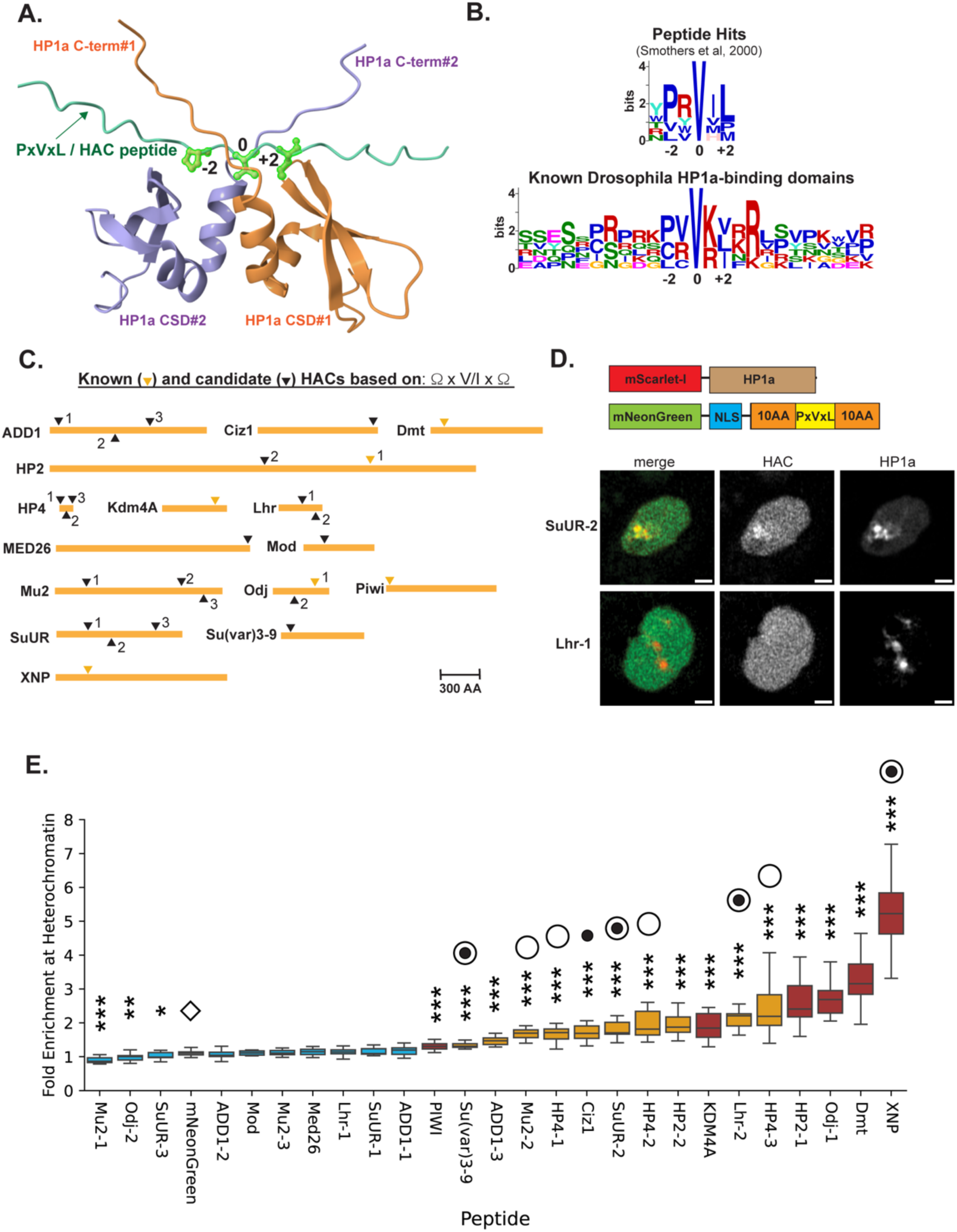
(A) Alphafold-generated model of a HP1a CSD dimer and a peptide containing the PxVxL motif. Sidechains of PxVxL positions-2, 0, and +2 are shown in green. (B) Motif analysis of original peptide sequences identified by phage display (Smothers et al, 2000, top) and the 6 sequences previously identified as HP1a-binding in *Drosophila melanogaster* (bottom). Core PxVxL residues-2, 0, and +2 are marked. (C) Schematic diagram of HP1a-binding proteins and their corresponding known (yellow triangles) or putative (black triangles) HACs. Numbers indicate positions of putative HACs on full-length proteins from N- to C- terminus, except for known HACs, which are automatically numbered 1. (D) Schematic of expression constructs mScarlet-I-tagged HP1a and mNeonGreen-tagged HACs (top) and representative images of constructs after transient transfection into cultured S2R+ cells (bottom). Scale bars = 1 um. (E) Fold enrichment of known and candidate HACs in heterochromatin are calculated by the ratio of mean intensity in heterochromatin to the mean intensity of the non-nucleolar nucleus. IDs of candidate HACs that fail to enrich in heterochromatin are highlighted in blue; known HACs are marked in red, and candidate HACs that display enrichment in heterochromatin are in orange. Also shown is the heterochromatin enrichment of the mNeonGreen tag, marked with a diamond, which equals about 1 and thus is used as the baseline to test significant enrichment of putative HACs by Student’s t-test. *** = p-value<0.001; ** = p-value<0.01; *=p-value<0.05. Determination of which candidate HACs are enriched in heterochromatin was also assessed manually as some candidates that did not visually display heterochromatin enrichment produced higher and lower enrichment values than the mNeonGreen control, likely due to some peptides accumulating at the nucleolus or being excluded from heterochromatin, respectively. HAC peptides tested for central residue mutation effects in this study are marked with dotted circles; HACs shown to be functional in the full-length protein are marked with empty circles.

HP1a binding sites have been mapped in many proteins, but of those only 6 in Drosophila have been shown to contain functional PxVxL motifs, as demonstrated by disrupting HP1a association through mutation of the central Val residue. These include the SNF2-like chromatin remodeler XNP (Bassett *et al*., 2008), histone H3 K36 demethylase KDM4A (Lin *et al*., 2008), zinc finger protein Odj (Kasinathan *et al*., 2020), structural heterochromatin protein HP2 (Stephens *et al*., 2005), sister chromatid cohesion regulator Dmt (Yamada *et al*., 2017), and transposon repressor Piwi (Brower-Toland *et al*., 2007) (**Supplementary Figure 1A**). A sequence alignment showed that while position 0 is consistently valine, the other positions vary between 3 hydrophobic residues (proline, leucine, and cysteine at-2 and leucine, isoleucine, valine at +2) (**Figure 1B**). Using the expanded consensus signature of HP1a-binding domains at positions-2 and +2, we probed the Drosophila proteome, as well as proteins annotated to localize to the nucleus or cytoplasm, for potential HP1a interactors (**Supplementary Figure 1B**). Not surprisingly, such a small motif identified over 50% of the proteome as HP1a interactors, including more than 50% of proteins annotated as cytoplasmic, where HP1a is not normally found. This led us to investigate whether the consensus sequence and our ability to predict HP1a binding domains can be enhanced. We note, for example, that positively charged residues frequently occupy the-1 and +1 positions along with flanking regions (**Figure 1B**), suggesting that other positions on the HP1a-binding sequence contribute to their function and could improve our understanding of the functional ‘HP1a Access Code’, hereafter referred to as “HAC.”

In order to discover additional HAC features, we needed to expand the number of sequences that exhibit HP1a association and differentiate them from those that lack such function. We assessed diverse HAC candidate sequences, surveying known HP1a-interacting proteins for a more degenerate pentamer sequence consisting of any hydrophobic amino acid at positions-2 and +2, and a central valine or isoleucine. The inclusion of isoleucine is based on sequence alignment of Dmt homologs, some of which were found to contain this residue instead of valine (**Supplementary Figure 2B**). We also focused our search on HP1a interactors that have been partially mapped for HP1a-interacting regions (Pindyurin *et al*., 2008; Schwendemann *et al*., 2008). Twenty putative HACs were identified in 11 proteins (**Figure 1C**), including HP2 and Odj that were already known to have at least one functional HAC. To test for HAC functionality, we employed an *in vivo*, imaging-based approach to screen 25-amino acid peptides from the HAC candidates for enrichment within the heterochromatin subnuclear compartment, as marked by fluorescently tagged HP1a. Similarly-sized peptides were previously shown to bind HP1a *in vitro* (Mendez *et al*., 2011), and the relatively short sequences minimize the possibility of including additional domains that may confound results. Thus, this assay indirectly tests HP1a binding through the ability of HAC-containing peptides to localize to the HP1a-rich heterochromatin, but also exposes candidate sequences to physiological conditions not possible with *in vitro* binding assays.

For our experiment, we tested peptides containing our known and candidate HACs, comprised of the hydrophobic pentameric core containing a PxVxL-like consensus plus 10-amino acid sequences on each flank, by fusing them to a fluorescent tag (mNeonGreen) and a myc nuclear localization signal. Co-transfection into Drosophila S2R+ cells with mScarlet-I-tagged HP1a showed co-enrichment of HP1a with known HACs, indicating that HAC peptides are sufficient to bind HP1a *in vivo* (**Supplementary Figure 1C**). In contrast, expression of the fluorescent tag and the NLS without any HAC peptide failed to concentrate at HP1a-rich nuclear regions. We also mutated the central HAC residue of several peptides, resulting in loss of enrichment in heterochromatin (**Supplementary Figure 1C**). Based on these results, we conclude that the peptide-based imaging approach is sufficient to detect sequences capable of associating with HP1a, as measured by fold-enrichment within the heterochromatin domain over the total nuclear signal.

Next, we tested putative HACs from several heterochromatin-interacting proteins and identified sequences from SuUR (Pindyurin *et al*., 2008), Lhr (Brideau and Barbash, 2011), HP4 (Schwendemann *et al*., 2008), Mu2 (Dronamraju and Mason, 2011), ADD1 (Alekseyenko *et al*., 2014), Ciz1 (Alekseyenko *et al*., 2014), HP2 (Stephens *et al*., 2005), Odj (Kasinathan *et al*., 2020), Mod (Perrin *et al*., 1998), MED26 (Marr *et al*., 2014), and Su(var)3-9 (Schotta *et al*., 2002) that did or did not accumulate in heterochromatin (**Figure 1D**). Heterochromatin-enriched peptides measured significantly higher levels of fluorescence intensity at regions of high HP1a enrichment relative to expression of the fluorescent tag without a HAC sequence (**Figure 1E**). This expands the number of known HACs from 6 to 16, while eliminating 10 sequences as putative HACs, or at the very least, HACs with HP1a affinities too low to detect with this assay. Image analysis also revealed varying levels of HAC co-enrichment with heterochromatin, with XNP displaying the highest co-enrichment with HP1a, and Piwi exhibiting the lowest. Mutation of the central residue of 4 of these peptides severely impaired heterochromatin localization, confirming that these comprise functional HAC sequences (**Supplementary Figure 1D**). We conclude that the *in vivo* peptide imaging approach can successfully differentiate varying levels of HP1a affinity among HACs, and that motifs resembling a PxVxL are frequently found in many proteins but poorly predict heterochromatin localization.

Finally, we determined whether the results of our peptide imaging experiment can be recapitulated with full-length proteins. Central residue mutations of HAC sequences in SuUR, Lhr and Mu2 abrogated enrichment to the heterochromatin domain and resulted in lower protein levels (**Supplementary Figure 1E-G**). This indicates that loss of binding with HP1a destabilizes these proteins, as has been shown for other complex-forming proteins (Hsu, Yen and Yeang, 2022). In contrast, corresponding mutations of non-functional sequences displayed no effect on heterochromatin co-localization or protein levels. Meanwhile, mutations in both the HAC and the H3K9me3-binding chromodomain (Wang *et al*., 2012) were necessary to abolish heterochromatin co-localization of Su(var)3-9 (**Supplementary Figure 1H**), highlighting how other domains also play roles in heterochromatin enrichment. Finally, HP4 required the simultaneous mutation of all 3 central HAC residues to fully abrogate the localization of HP4 to heterochromatin (**Supplementary Figure 1I**), confirming their redundant functions to target HP1a. We therefore conclude that new HACs identified by peptide localization correspond to functional heterochromatin localization domains in full-length proteins.

### Drosophila HACs are conserved yet embedded in disordered regions

In order to further define the sequence features that mediate HP1a binding and heterochromatin-targeting, we investigated whether functional HACs are more evolutionarily-conserved compared to non-functional peptides. Sequence alignment of HP1a CSD and C-terminal tails for 12 Drosophila species across 40 million years of evolution show remarkable conservation (91-100% identity in the CSD, 100% in the C-terminal tail) (**Supplementary Figure 2A**). Sequence alignments of heterochromatin proteins harboring previously known HACs and their homologs also show a high degree of conservation in terms of containing a degenerate HAC pentamer sequence at the same position in the protein (**Figure 2A left, Supplementary Figure 2B**). Kdm4A, Dmt, HP2, and Odj HACs are preserved across all 12 Drosophila species, though the Piwi HAC appears to be more recently evolved. Similar results were obtained with our newly identified HACs, with 6 of 10 conserved across Drosophilids (**Figure 2A middle, Supplementary Figure 2B**). In contrast, the non-functional candidate sequences, as a group, were generally less conserved compared to functional ones (mean conservation = 70% vs 87%) (**Figure 2A right, Supplementary Figure 2B**). However, the non-functional candidate sequences with high conservation could be part of other domains.

**Figure 2.**
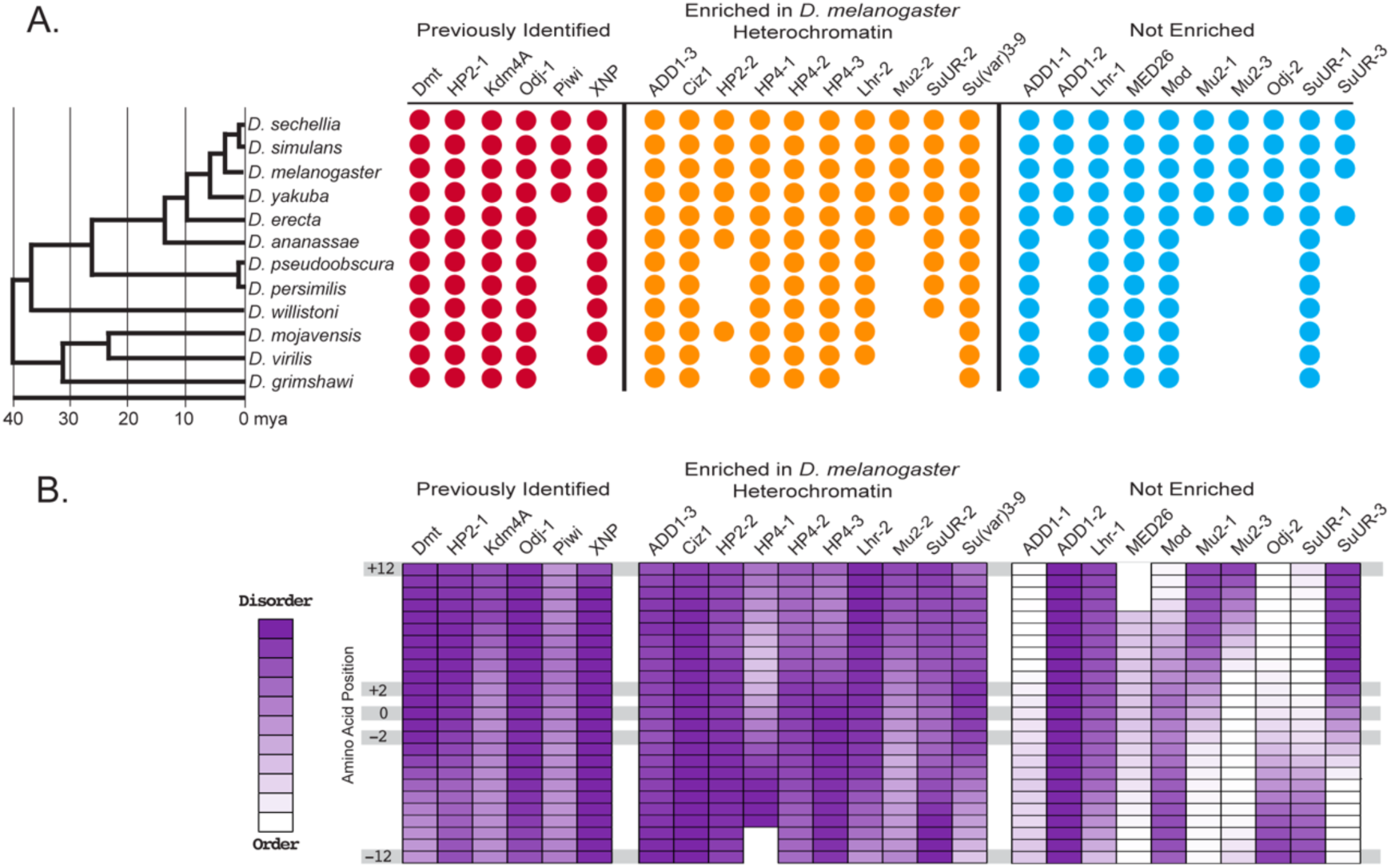
(A) Conservation of HAC sequences tested in this study among Drosophila species spanning up to 40 million years of evolution (tree indicated on the left). Positive identification was assessed by presence of [hydrophobic]x[V/I]x[hydrophobic] pattern after alignment to *D. melanogaster* sequences among previously known HACs (red circles), as well as positively identified HACs (yellow circles) and sequences that did not display HP1a co-enrichment (blue circles) from Figure 1E. (B) Heat map showing disorder prediction per residue of HAC sequences in (A) as obtained from the PONDR-VLXT algorithm. Prediction scores indicate presence of intrinsic disorder (purple) to globular structures (white). Core HAC residues at - 12,-2, 0, +2, and +12 positions are indicated.

The higher degree of conservation among functional HAC sequences suggests that they may also share an ordered structure. However, disorder prediction analysis of the entire protein sequence using PONDR (Romero *et al*., 2001) showed that known HACs are either embedded within disordered domains, or in transition regions between ordered and disordered regions (**Figure 2B left**). Similarly, newly identified HACs showed a similar association with disordered regions (**Figure 2B middle**). In contrast, the group of non-functional sequences included both disordered and highly ordered structures (**Figure 2B right**). This indicates that functional HACs, despite their localization in disordered regions, which are often associated with higher mutation rates than globular domains, may be subjected to higher purifying selection for conservation. This also suggests that HACs inherently occur in the more solvent-accessible unstructured regions of proteins to facilitate binding to the HP1a CSD dimer. Overall, our results indicate that high evolutionary conservation and association with disordered domains are two shared features of HACs.

### The pentameric core requires flanking residues for heterochromatin localization

To identify the amino acids and chemical interactions responsible for binding of HACs with HP1a dimers, we performed full atomistic molecular dynamics (MD) simulations on 25-amino acid HAC sequences containing the hydrophobic core at the center. We confirmed that 15 of the 16 HACs showed high prediction accuracy with pLDDT (predicted Local Distance Difference Test) (**Supplementary Method Table 1**), and the expected positioning of their core region at the center of the CSD dimer in AlphaFold models and throughout the 250 ns MD simulations (**Figure 3A, Supplementary Figure 3A-O**). To investigate whether the core region is sufficient to explain the variable fold-enrichment of HACs in heterochromatin (**Figure 1E**), we compared the root mean square deviation (RMSD) of HAC sequences from - 2 to +2 in the last 50 ns of the MD simulations (**Figure 3B)**. Though Piwi, with the lowest fold enrichment score, showed very high fluctuation, the RMSD of the core regions of other peptides showed poor correlation with the ranking of HP1a co-enrichment scores from experimental observations (**Figures 1E, 3B,** R^2^ = 0.07). To expand and elucidate key residues of HACs, the root mean square fluctuation (RMSF) was calculated for Piwi and XNP, which had the lowest and highest fold enrichment scores from the peptide imaging assay, respectively (**Figure 3C**). XNP showed overall lower RMSF values than Piwi as well as a longer region of low RMSF values (≤RMSF values of the Piwi core region, dotted horizontal lines, **Figure 3C**), consistent with the stronger enrichment of the XNP HAC with HP1a in cells (**Figure 1E**). As expected, the core regions (-2 to +2) exhibited the lowest RMSF values, indicating the part of the HAC with the most stable association with the CSD binding pocket (**Figure 3C**). Interestingly, the proximal flanking sequences on both sides (-7 to-3 and +3 to +7) also showed low RMSF values and no significant difference from the core regions of both XNP and Piwi (**Figure 3C**). While Piwi showed no significant RMSF difference between the proximal flanking sequences and distal flanking sequences (-12 to-8 and +8 to +12), XNP had a significant difference between the proximal and distal flanking sequences (**Figure 3C**). These RMSF results from the MD simulations suggest the potentially crucial role of the proximal flanking regions in determining the strength of the HP1a-peptide interactions and accessibility to heterochromatin.

**Figure 3.**
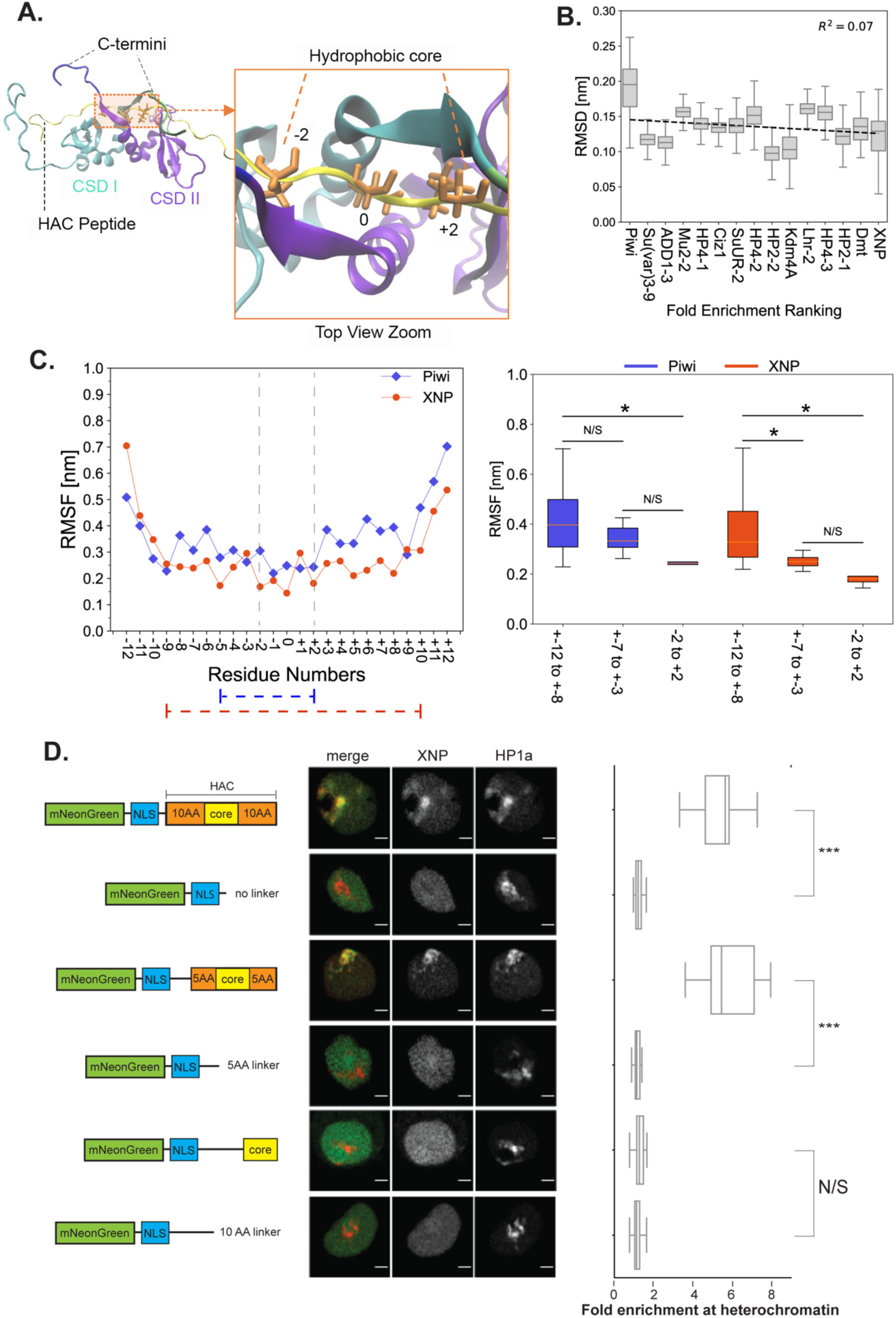
(A) Visualization of the CSD dimer with the XNP HAC. Left panel: side view. Right Panel: Zoomed top view. Each CSD monomer is colored cyan and purple with a green and blue colored C-terminal tail, respectively. XNP peptide is colored yellow. Residues of the central hydrophobic core at the-2, 0, and +2 positions (in licorice drawing style) are colored orange. (B) Root mean square deviation (RMSD) ± SD [nm] of the core sequences from-2 to +2 of 15 HACs between 200 to 250 ns of the MD simulations. The peptides are ranked according to the experimental ranking of fold HP1a co-enrichment from Piwi, the lowest, to XNP, the strongest, based on results from Figure 1E. The x-axis values for the linear regression calculation are assigned from 1 (Piwi) to 15 (XNP). The R-squared value is shown on the graph. (C) Left: Root mean square fluctuation (RMSF) [nm] of 25-residue for XNP and Piwi HAC sequences (-12 to +12) between 200 to 250 ns of the MD simulations. The gray dash line at the-2 and +2 is drawn to indicate the core pentamer. The blue and red dashed lines indicate the span of sequences that consecutively display RMSF values less than the combined average of Piwi and XNP = 0.32 nm. Right: Averaged root mean square fluctuation (RMSF) ± SD [nm] of the XNP and Piwi distal flanking sequences (positions-12 to-8 and +8 to +12), proximal flanking sequences (positions-7 to-3 and +3 to +7), and core sequences (positions-2 to +2). Asterisks denote p-values < 0.05 (*) (D) Left: Representative localization images of XNP HAC constructs with different amounts of flanking residues around the hydrophobic core as illustrated in the right panel. As controls, fluorescent tags with different lengths of linker regions were also tested. Right: Quantitation of HP1a co-enrichment at heterochromatin domains for the tested HAC constructs. N/S = not significant; *** = p-value < 0.001.

We tested different lengths of flanking sequences required for the central core of HACs to maintain heterochromatin enrichment *in vivo*. Lack of any flanking sequences abrogated the ability of three different HACs to localize to heterochromatin at levels indistinguishable from negative controls (**Figure 3D**, **Supplementary Figure 3P and Q**). In contrast, the retention of 5 amino acids flanking each side of the central pentamer maintained enrichment with HP1a, indicating that flanking residues are crucial to HAC association with HP1a (**Figure 3D, Supplementary Figure 3P**). In conclusion, both MD simulations and *in vivo* peptide localization demonstrate that the traditional PxVxL motif is insufficient for heterochromatin enrichment, and additional residues within 5 amino acids on both sides of the hydrophobic core are required for HAC function.

### Electrostatic forces surrounding the hydrophobic pocket drive HAC enrichment to heterochromatin

To identify the driving force of the interactions between HACs and HP1a, the types of amino acids in HACs and their molecular properties were analyzed *in silico*. Sequence alignment of all 16 HACs displayed high frequencies of hydrophobic amino acids at positions-7,-5,-1, +5, and +7 in addition to the core hydrophobic residues (-2, 0, and +2) (**Figure 4A**). Amino acids with positively charged side chains (Lys and Arg) are enriched in the flanks and interior of the central hydrophobic core (-4,-3, +1, +3, and +4) when compared to non-HAC peptides (**Figure 4A, Supplementary Figure 4A**). In contrast, negatively charged residues (Asp and Glu) are relatively under-represented among HAC peptides. The sequence alignment analysis thus suggests that hydrophobic and positively charged amino acids contribute significantly to HP1a interactions at specific positions of HACs.

**Figure 4.**
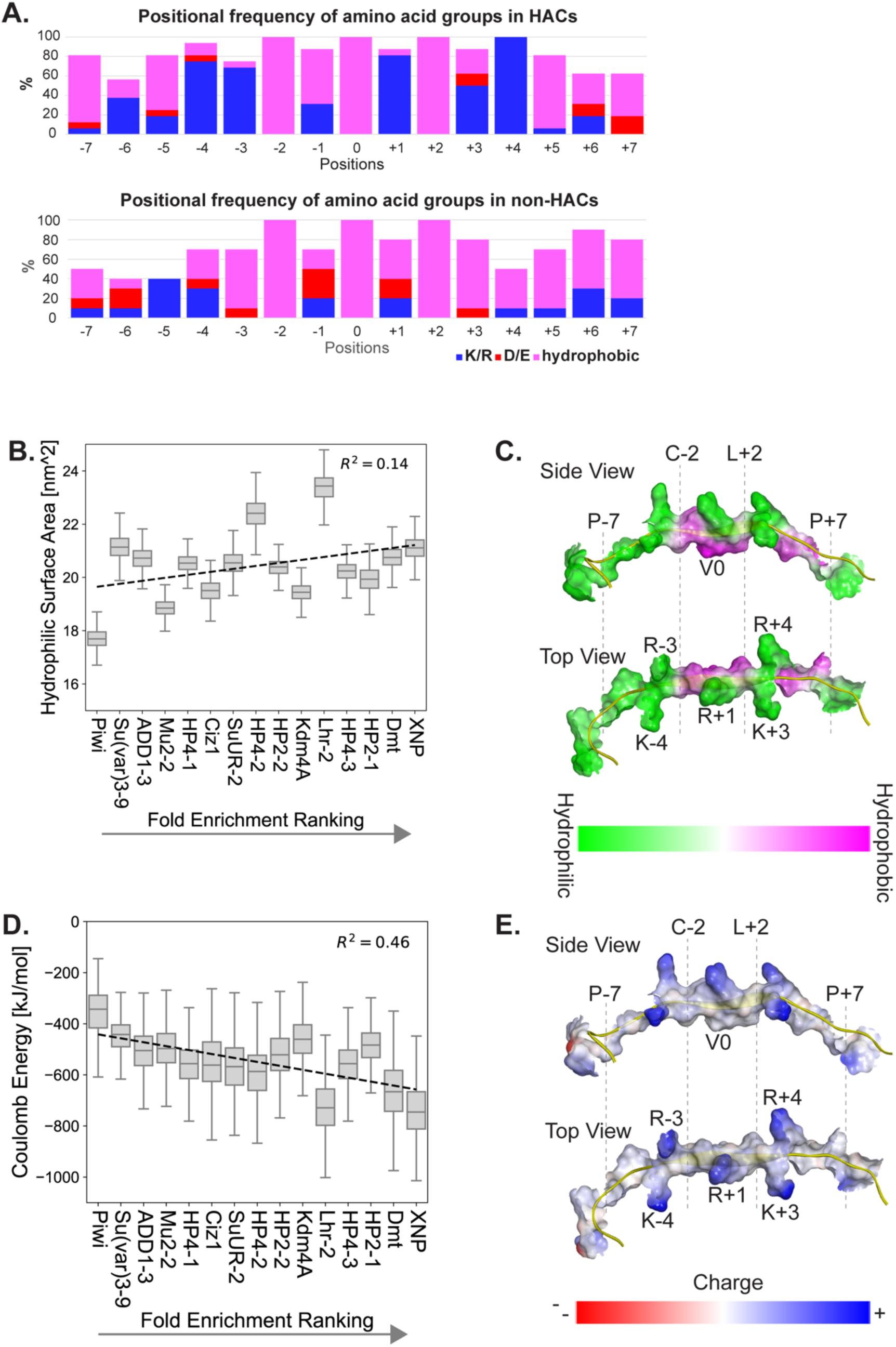
(A) Frequency of occupancy by hydrophobic (pink), positively charged (blue), and negatively charged (red) select groups of amino acids at each position among tested HACs (top) and non-HACs (bottom). Numbering of positions are in relation to the central residue at position 0. (B) Hydrophilic surface area ± SD [nm^2^] of sequences from-7 to +7 of 15 HACs between 200 to 250 ns of the MD simulations. The peptides are ranked according to the experimental ranking of fold HP1a co-enrichment from Piwi, the lowest peptide, to XNP, the strongest peptide. The x-axis values for the linear regression calculation are assigned from 1 (Piwi) to 15 (XNP). The R-squared value is shown. (C) Visualizations of XNP HAC surface hydrophilicity and hydrophobicity in the side view (above) and the top view (bottom) were calculated using the Discovery Studio software. The hydrophilic to hydrophobic properties are colored from green to pink. XNP backbone is colored yellow. (D) Coulomb energy ± SD [kJ/mol] of 15 HAC sequences from-4 to +4 between 200 to 250 ns of the MD simulations. A ranking similar to that of (A) is used to order the peptides. The R-squared value is shown. (E) Visualization of the XNP HAC surface charge in the side view (above) and the top view (bottom). The negatively to positively charged properties are colored from red to blue. XNP is colored yellow.

We first investigated the effects of the hydrophobic properties of HACs using MD simulations. The hydrophobic pocket formed by the central pentamer of HACs and the HP1 CSD dimer was previously identified to be the main interaction region. The flanking hydrophobic residues, such as-5 and +5, were also thought to contribute to hydrophobic interactions (Thiru *et al*., 2004). We compared the hydrophilic surface area of the HAC sequences from-7 to +7 obtained from the MD analysis with the experimental ranking of HP1a co-enrichment (**Figure 4B, 1E**). Surprisingly, despite some variations, the hydrophilic surface area values did not correlate with the fold enrichment ranking (**Figure 4B, Supplementary Figure 4B,** R^2^ = 0.14). The hydrophilic surface area of only the HAC central pentamer from-2 to +2 also showed no correlation with the fold enrichment scores (**Supplementary Figure 4D,** R^2^ = 0). Comparing the hydrophilic surface area frequencies of the top two and bottom two ranking peptides showed that Piwi had the highest and lowest hydrophilic surface area in the HAC core pentamer (-2 to +2) and the proximal flanking region (-7 to +7), respectively (**Supplementary Figure 4C, E**). This indicates that Piwi possesses the lowest hydrophobic interaction between the CSD dimer and the HAC core pentamer, but the highest hydrophobicity between the CSD and the proximal flanking sequence (**Supplementary Figure 4C, E**). However, Su(var)3-9 and Dmt, which had the second lowest and second highest fold enrichment scores, respectively, did not show a corresponding trend of hydrophilic surface area values in relation to Piwi and XNP (**Supplementary Figure 4C, E**).

To investigate how the hydrophilicity and hydrophobicity of each residue on the HAC peptide participate in the interaction, we visualized the hydrophilic and hydrophobic distribution properties of the HP1a-interacting XNP regions (**Figure 4C**). As expected, there is a hydrophobic pocket between the-2 to +2 amino acid sequences of the HAC and HP1 CSD dimer (**Figure 4C**). On the contrary, the nearby flanking residues tend to have high hydrophilicity rather than hydrophobicity (**Figure 4C**). The hydrophobic pocket of the peptide and α-helices of the CSD dimer were surrounded by a high hydrophilic region composed of - 4,-3, +1, +3, and +4 positions, indicating the possible existence of other types of interactions that could modulate interactional strength with HP1a. These results also suggest that the hydrophobic interactions of the HACs alone are insufficient to describe the fold enrichment of HAC-containing peptides with HP1a.

Given the lack of correlation between the hydrophobic properties and strength of HP1a co-localization *in vivo*, we then investigated the electrostatic contributions of the core and flanking amino acids using MD simulations. Electrostatic interactions, which include salt bridges, pi-cation interactions, and hydrogen bonding, represent another key type of interaction in protein structure and binding (Zhou and Pang, 2018). Salt bridge interactions, in particular, are known to be one of the most robust forms of residue-residue interactions (Xie *et al*., 2015). Salt bridge interactions form when the distance between the side-chain carbonyl oxygen of Asp or Glu and the side-chain nitrogen atoms of Arg, Lys, or His is within 0.4 nm, facilitated by Coulomb attraction with water molecules (Kumar and Nussinov, 2002; Pylaeva, Brehm and Sebastiani, 2018). The high hydrophilic and positively charged amino acid-conserved regions were seen at-4,-3, +1, +3, and +4 (**Figure 4A, C**). Therefore, we calculated Coulomb energy levels between the HACs from +4 to +4 and the HP1a CSD dimer with C-terminal tails. Notably, Coulomb energy levels generally decreased among peptides ranked from low to high HP1a co-enrichment, indicating that attractive electrostatic force followed the trend of the *in vivo* fold enrichment ranking (**Figure 4D, Supplementary Figure F**, R^2^ = 0.46). Comparing the electrostatic energy of the top two and bottom two ranking peptides similarly showed a correlation between Coulomb energy levels and heterochromatin enrichment (**Supplementary Figure G**). The Coulomb energy between the HACs from-7 to +7 and the HP1a CSD dimer with C-terminal tails showed less correlation (**Supplementary Figure H, I**, R^2^ = 0.28), implying the-4 to +4 residues of HACs have more important contributions to the fold enrichment scores. The visualization of the electrical charge distribution of the HP1a-interacting XNP regions showed a high charge density at-4,-3, +1, +3, and +4 amino acids outside the hydrophobic pocket of the CSD, but did not show a charge density outside-4 and +4 regions, emphasizing the importance of electrically charged amino acids between-4 and +4 in the electrostatic interactions with HP1a (**Figure 4E**).

Overall, our observations indicate that electrostatic interactions between the HAC residues from-4 to +4 and the HP1a CSD dimer with C-terminal tails are crucial determinants in the binding of HAC-containing peptides with HP1a dimers. Of course, hydrophobic and electrostatic residues beyond the ranges of positions analyzed here may also contribute to HP1a binding, but these positions, such as distant flanking sequences (-12 to-8 and +8 to +12), are more variable in residue content and experience more fluctuation (as shown by RMSF variation in Figure 3).

### Distinct roles of positively charged amino acids in HAC binding

To reveal the molecular roles of positively charged amino acids in the HAC peptide binding and structure, we analyzed the individual Coulomb energy from positions-7 to +7 for several HACs in association with the HP1a CSD dimer and C-termini in the MD simulations (**Figure 5A, C**). Three peptides were chosen for the electrostatic analysis: XNP as the strongest interacting peptide, Piwi as the weakest interacting peptide lacking a positively charged amino acid at the +3 position, and Ciz1 as the relatively weak interacting peptide lacking a positive amino acid at the +1 position. **Table 1** shows the quantitative values of Coulomb energy for - 7 to +7 sequences of these three peptides. Overall,-4,-3, +3, +4 HAC amino acids had higher attractive Coulomb energy than-7 to-5 and +5 to +7 regions, quantitatively confirming the important contributions of-4 to +4 amino acids suggested in Figure 4.

**Figure 5:**
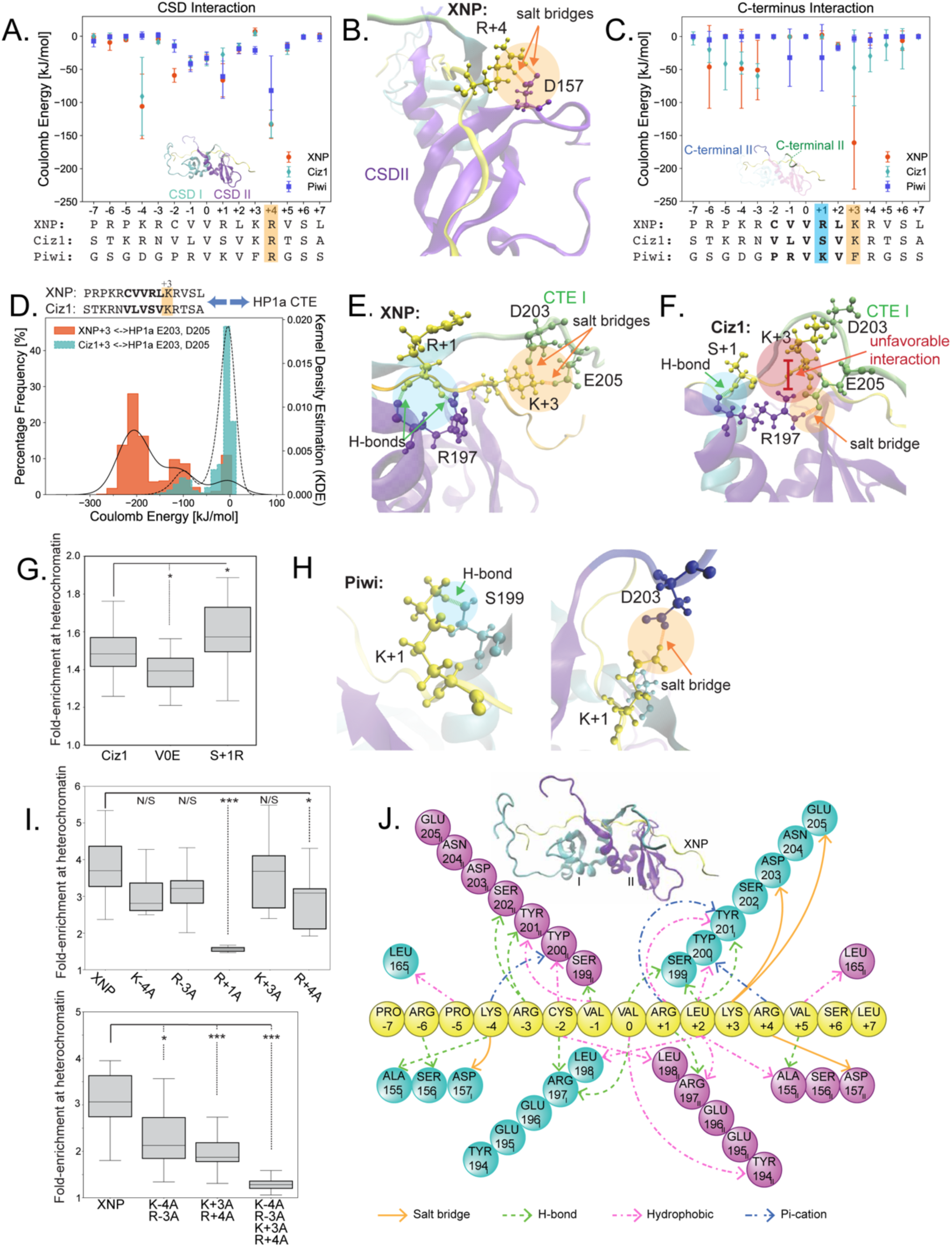
(A) Coulomb energy ± SD [kJ/mol] between the CSD dimer and-7 to +7 residues of XNP, Ciz1, and Piwi. Orange circle: XNP, Cyan diamond: Ciz1, and Blue square: Piwi. (B) Visualization of the salt bridge interaction between Arg+4 and Asp157^II^. Arg+4 and Asp157^II^ in the Corey-Pauling-Koltun (CPK) drawing style are colored yellow and purple, respectively. (B) Coulomb energy ± SD [kJ/mol] between the HP1a C-terminal tails and-7 to +7 residues of XNP, Ciz1, and Piwi. Orange circle: XNP, Cyan diamond: Ciz1, and Blue square: Piwi. (D) Histograms showing the percentage frequency of Coulomb energy between HP1a Asp203^I^/Glu205^I^ and +3 of XNP and Ciz1 between 200 to 250 ns of the MD simulations. The histogram was fitted by kernel density estimation (KDE). XNP: Orange with a solid line. Ciz1: Cyan with a dashed line. (E) Visualizations of XNP and (F) Ciz1 with blue, orange, and red circles with arrows indicating a hydrogen bond, salt bridge, and unfavorable interactions, respectively. (G) Quantification of HP1a co-enrichment of Ciz1 HAC and mutations either disabling the central valine or adding a positive charge at the +1 position. (H) Visualization of H-bond interaction between Piwi Lys+1 and Ser199^I^ (left) and salt bridge interaction between Piwi Lys+1 and Asp203^II^ (right). H-bond and salt bridge are highlighted in green and orange, respectively. (I) Quantification of HP1a co-enrichment of XNP HACs with single alanine substitutions at positions-4,-3, +1, +3, or +4 (top) or double substitutions at positions-4 and-3 or +3 and +4 or quadruple substitutions at all positions (bottom). Asterisks denote p-values < 0.05 (*), <0.01 (**), <0.001 (***). (J) 2D molecular interactional map of HP1a CSD dimer with C-terminal tails and XNP HAC. Each CSD monomer is colored cyan and purple, respectively. XNP is colored yellow. Orange, green, pink, and blue dashed lines represent a salt bridge interaction, hydrogen bond, hydrophobic interaction, and pi-cation interaction.

**Table 1.** Sequences and coulomb energy ± SD [kJ/mol] among the CSD dimer or CTE and-7 to +7 residues of XNP, Ciz1, and Piwi. The highest coulomb energy among the-7 to +7 is highlighted in red.

| Residue | XNP | Ciz1 | Piwi | CSD_XNP | CSD_Ciz1 | CSD_Piwi | C-terminus_XNP | C-terminus_Ciz1 | C-terminus_Piwi |
| --- | --- | --- | --- | --- | --- | --- | --- | --- | --- |
| -7 | P | S | G | -1.02 ± 4.99 | 0.17 ± 0.89 | -7.34 ± 8.71 | -0.04 ± 0.13 | -2.79 ± 9.95 | 0.02 ± 2.09 |
| -6 | R | T | S | -9.27 ± 11.92 | -1.94 ± 6.99 | -0.89 ± 2.73 | -46.09 ± 62.83 | -20.22 ± 19.17 | -5.35 ± 13.21 |
| -5 | P | K | G | -5.45 ± 2.38 | -2.55 ± 4.81 | 0.09 ± 0.55 | 0.60 ± 1.37 | -41.43 ± 39.44 | -0.10 ± 1.04 |
| -4 | K | R | D | -105.93 ± 48.80 | -90.84 ± 59.05 | 1.06 ± 5.62 | -49.02 ± 59.89 | -40.17 ± 16.80 | 0.00 ± 7.94 |
| -3 | R | N | G | -4.58 ± 5.44 | -8.29 ± 2.93 | 1.32 ± 4.32 | -50.86 ± 45.06 | -59.87 ± 18.65 | 0.23 ± 0.22 |
| -2 | C | V | P | -59.14 ± 10.41 | -14.00 ± 3.01 | -14.79 ± 9.36 | -0.84 ± 1.17 | -0.58 ± 2.31 | -0.01 ± 0.05 |
| -1 | V | L | R | -41.48 ± 9.21 | -38.41 ± 7.45 | -41.34 ± 12.06 | -0.35 ± 0.75 | -0.33 ± 0.62 | -32.02 ± 43.60 |
| 0 | V | V | V | -34.87 ± 7.38 | -34.82 ± 9.73 | -32.84 ± 9.36 | -0.13 ± 0.27 | -0.02 ± 0.03 | 0.04 ± 0.05 |
| +1 | R | S | K | -66.45 ± 25.07 | -27.53 ± 9.84 | -60.91 ± 32.91 | 2.45 ± 6.20 | -0.19 ± 1.72 | -32.08 ± 50.42 |
| +2 | L | V | V | -20.24 ± 5.45 | -15.57 ± 5.31 | -18.88 ± 7.60 | -13.30 ± 4.62 | -14.29 ± 4.19 | -16.89 ± 4.62 |
| +3 | K | K | F | 7.16 ± 5.26 | 4.82 ± 4.27 | -21.33 ± 5.79 | -161.04 ± 70.42 | -47.46 ± 57.53 | -3.01 ± 5.15 |
| +4 | R | R | R | -133.15 ± 21.40 | -132.20 ± 21.07 | -81.99 ± 52.34 | -3.96 ± 9.15 | -29.54 ± 23.95 | -6.04 ± 11.63 |
| +5 | V | T | G | -16.68 ± 5.73 | -18.45 ± 8.98 | -15.14 ± 6.16 | -0.05 ± 1.17 | -12.96 ± 23.31 | 0.04 ± 0.38 |
| +6 | S | S | S | -0.93 ± 1.01 | -0.69 ± 2.24 | -1.21 ± 3.60 | -6.33 ± 14.80 | -18.59 ± 30.59 | -0.11 ± 0.86 |
| +7 | L | A | S | 1.49 ± 0.99 | -0.37 ± 2.36 | -1.34 ± 6.45 | 0.83 ± 2.43 | -0.28 ± 1.13 | -0.01 ± 0.23 |

The highest attractive energy to the CSD dimer was observed at the +4 position for all three peptides (**Table 1**, **Figure 5A**), where a salt bridge interaction with Asp157^II^ (of CSD^II^) was detected (**Figure 5B, Supplementary Figure 5A**). Similarly, the-4 position of the XNP and Ciz1 HACs formed a salt bridge interaction with Asp157^I^, but was not present for Piwi due to the lack of a positively charged amino acid at the-4 position (**Figure 5A**). All 16 peptides that exhibited heterochromatin enrichment have a positively charged amino acid at the +4 position (**Figure 4A**), suggesting the +4 positively charged amino acid comprises a basic feature of the HAC motif.

In contrast, the residues with the highest attractive Coulomb energy with the HP1a C-terminus varied among the three peptides (**Table 1**). The Coulomb energy at the +3 position displayed high variance among the three peptides (-164±70.42,-47.39±57.50, and-3.01±5.15 [kJ/mol] for XNP, Ciz1, and Piwi, respectively). Unlike the other two, Piwi lacks a positively charged amino acid at the +3 position. Interestingly, there is also a discrepancy between XNP and Ciz1, even though both XNP and Ciz1 have lysine at the +3 position. Molecular interaction analysis showed that the Arg+3 of XNP formed a salt bridge or H-bond with HP1a Glu205^I^ or Asp203^I^ (**Figure 5D, Supplementary Figure 5B**). However, Arg+3 of Ciz1 did not form such interactions and instead had an unfavorable interaction with HP1a Arg197^II^ (**Figure 5D, F, Supplementary Figure 5C**). This difference is attributed to interactions between +1 of the HACs and Arg197^II^ of the CSD: Arg+1 of XNP formed a maximum of three H-bonds with Arg197^II^, including its side chain (**Figure 5E, Supplementary 5C**), while Ser+1 of Ciz1 had only one H-bond with the carboxylic acid group of Arg197^II^, leading to a repulsive interaction between the Arg197^II^ side chains and Lys+3, and a H-bond between Arg197^II^ and Glu205^I^ (**Figure 5F, Supplementary 5C**). These results indicate that Arg+1 plays an important role in both the H-bond interaction with Arg197^II^ and the maintenance of the electrostatic force field to develop the salt bridge and H-bond among +3, Asp203^I^, and Glu203^I^. We therefore tested the effects of a Ser+1Arg Ciz1 mutation *in vivo* to determine if increasing electrostatic interactions at this position will enhance HP1a co-enrichment. The Ciz1 mutant modestly improved the fold enrichment score compared to a wildtype Ciz1 HAC, indicating the deterministic effect of +1 in controlling the strength of the HP1a and HAC peptide interaction (**Figure 5G, Supplementary Figure 5D**).

As previously noted, Piwi lacks a positively charged amino acid at +3, but our analysis showed that the Lys+1 side chain can form an H-bond with Ser199^I^ or salt bridge interaction with Asp203^II^ (**Figure 5H, Supplementary Figure 5E**). A positively charged amino acid at position +1 may therefore compensate for the lack of a positively charge at +3 by forming a salt bridge or hydrogen bond to attain sufficient HP1a interaction strength. There are likely many other compensatory mechanisms for different HACs, as positively charged residues occupy variable positions within and surrounding the hydrophobic core (**Supplementary Figure 5F**).

To validate the computational theory of the electrostatic effects of positive amino acids on HAC peptide and HP1 interactions, we performed single and combinatorial amino acid mutation experiments with XNP, which had the highest Coulomb energy for the C-terminus and CSD dimer (**Table 1**, **Figure 5A, C**). First, single mutation experiments showed that Arg+1Ala and Arg+4Ala mutations significantly decreased the fold enrichment in heterochromatin (**Figure 5I, Supplementary Figure 5G**). Although high electrostatic energy was observed for Arg+3 in the MD simulation, the Arg+3Ala mutation did not show a significant difference in HP1a co-enrichment. The compensatory mechanism of the Piwi Arg+1 side chain for the lack of a positively charged amino acid at +3 (**Figure 5H**) might explain the lack of effect of Arg+3Ala in the XNP experiment. Nevertheless, the double mutations of Lys+3 and Arg+4, as well as double mutations of Lys-4 and Arg-3, greatly diminished XNP HAC enrichment in heterochromatin compared to the single mutants. Furthermore, a quadruple mutation of-4,-3, +3, and +4 nearly abrogated all enrichment, suggesting the basic residues act cooperatively to stabilize HAC enrichment to heterochromatin (**Figure 5I, Supplementary Figure 5H**). These results show that positively charged amino acids immediately flanking the pentameric core are essential for HAC function. Further, they suggest that the interaction strength between the HAC and HP1a can be controlled by selecting appropriate mutations of positively charged amino acids in the HAC sequence.

To obtain a comprehensive view of the molecular forces that govern HAC binding to HP1a, we analyzed the XNP HAC for electrostatic and hydrophobic interactions in simulations (**Figure 5J, Supplementary Figure 5I**). Limiting the scope of analysis to between HAC residues-7 and +7, we identified salt bridges forming at positions-4, +3, and +4 with both the CSD and C-terminal tail, as previously discussed. We also detected pi-cation interactions at the basic positions-4, +1, and +4 with the hydrophobic pocket residues of the CSD dimer. Meanwhile, hydrophobic interactions were predicted at positions-5,-2,-1, 0, +2, and +5, consistent with previous studies. Hydrogen bonds were also formed by both nonpolar and basic amino acids at 9 of the 15 residues analyzed. Of important note is the hydrogen bonding detected at position-2 unique to cysteine, due to its polar, hydrophobic sidechain. This confers an extra electrostatic interaction absent from other hydrophobic residues that occupy that position in other HACs, and thus provides higher affinity for HP1a. We predict that other HACs follow a general pattern of molecular interactions contained in this XNP map, though alternative interactions are likely to occur due to the positional variations of basic and hydrophobic residues of other HACs that impact HP1a affinity.

### Generation of an improved HAC molecular signature using homolog analysis

Our discovery of new HAC sequences, coupled with their high conservation between Drosophila species, amount to 173 peptide sequences that could be used to improve our understanding of the HAC molecular signature. Color-coded sequence alignment stacks of HACs and their homologs between hydrophobic, positively charged, and negatively charged residues reveal high conservation within each group of HAC homologs (**Figure 6A** and **Supplementary Figure 6A**). This suggests that HAC organization for each protein possesses specific features that contribute to its affinity for HP1a. For example, HACs containing a proline at-2 position often lack positively charged residues at the-3 and-4 positions, which may indicate a proline-imparted kinked structure that prohibits positively charged residues at those positions or render them unnecessary. We also identified rare, new residues at key positions such as leucine or asparagine at position +4, of *D. willistoni* HP2-1 and Mu2-3, respectively, capable of supporting HP1a co-enrichment (**Supplementary Figure 6B**), despite the other 171 sequences containing positively charged residues at that position.

**Figure 6.**
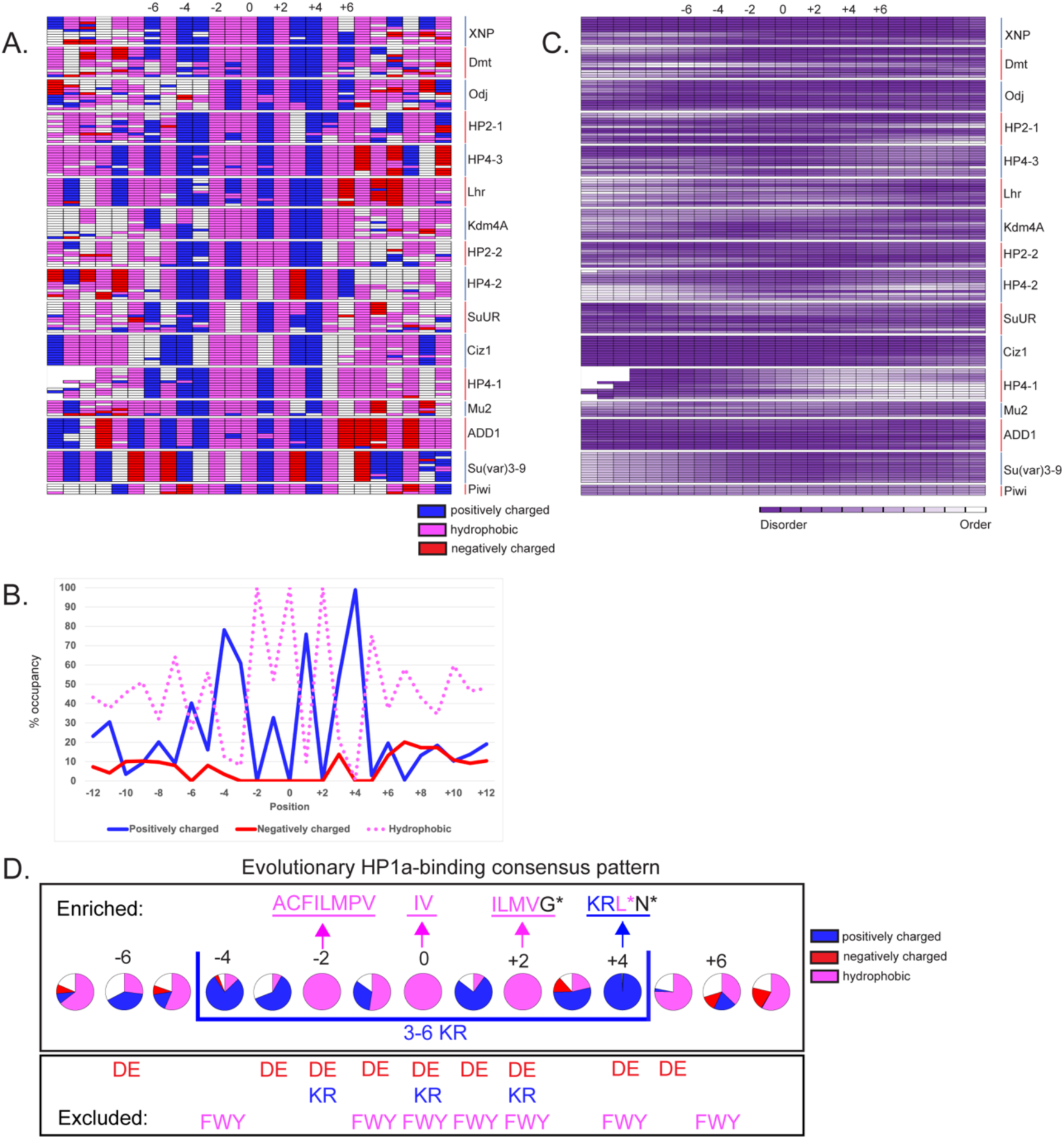
(A) Compilation of alignments for each HAC homolog across Drosophila evolution showing positioning of basic (red), hydrophobic (blue), and acidic (green) residues. Position numbers are indicated on the top and gene names are provided on the right of each stack of alignments. (B) Percent occupancy of basic (red), hydrophobic (dotted), and acidic (green) residues is calculated across Drosophila evolution for each position in all HACs. (C) Compilation of disorder prediction scores are shown for each position across evolution for each HAC. (D) Diagram of the evolutionarily conserved HAC molecular signature shows amino acids compatible for specific positions (top row), incompatible amino acids (second row), sidechain group bias (third row).

To identify unifying features of HACs, the occupancy of the same groups of amino acids were quantified for each position (**Figure 6B**). Aside from the expected 100% occupancy of nonpolar residues at positions-2, 0, and +2, there is nearly 80% occupancy at position +5 and about 60% at positions-7,-5,-1, and +7. This indicates that hydrophobic forces in the flanking regions of some HAC pentamers also contribute to functional HP1a binding directly or indirectly. Consistent with our analysis in previous figures, positively charged residues predominate at positions-6,-4,-3, +1, +3, and +4, which also correspond to a dearth of negatively charged residues. We hypothesize that the less frequent but conserved occupations of negatively charged residues in the flanking regions among specific HAC homologs reflect mechanisms to partially destabilize HP1a affinity and/or utilize alternative mechanisms of binding HP1a at those positions. This domination of HAC organization by nonpolar and basic residues is highlighted by the corresponding dramatic drop of amino acid heterogeneity between positions-6 and +6 (**Supplementary Figure 6C**). Finally, HAC organization across various Drosophila species is also marked by association with intrinsically disordered regions (**Figure 6C**), indicating that properties of these regions contribute to HAC function or that HAC sequences impart disorder to the surrounding protein structure.

To summarize our findings, conservation analysis of Drosophila homolog sequences revealed evolutionary positional bias for nonpolar and basic residues that correspond to various molecular interactions with the HP1a dimer (**Figure 5I**). We refine the original Drosophila HAC signature to consist of nearly any hydrophobic residue at-2, an invariant valine or isoleucine at the central position, and nearly any aliphatic amino acid at +2. Expanding this molecular signature, we also identify a nearly invariant basic residue at +4 (**Figure 6D**). We also observe basic residues to be favored at-4,-3, +1, and +3, while hydrophobic interactions are favored at positions-7,-5,-1, +5, and +7. However, due to the relative heterogeneity of residues found at these positions, they likely modulate HP1a interactions, as we showed previously for positions +1 and +3. Instead, we propose an enrichment of 3-6 basic residues between positions-4 to +4 as a key signature of HACs (**Figure 6D**). Additionally, no acidic residues are found at-6,-3,-1, +1 and +5, and no aromatic residues are located at-4 and +6, suggesting these are detrimental to HAC function.

Previously, we showed that the PxVxL motif based on known HP1a-binding sites is found in nearly half of all proteins, including those annotated to be in the nucleus or chromosomes **(Supplementary Figure 1B)**. In contrast, the updated HAC molecular signature reduces the number of potential candidates over 10-fold, even without accounting for disorder content **(Figure 7A)**. GO analysis of the proteins that contain sequences matching the updated HAC signature yielded heterochromatin and chromocenter as the top annotations (**Figure 7B**). In contrast, these categories were not significantly represented among the protein hits for the PxVxL motif search, indicating that our updated HAC signature vastly improves the prediction of HP1a-binding motifs.

**Figure 7:**
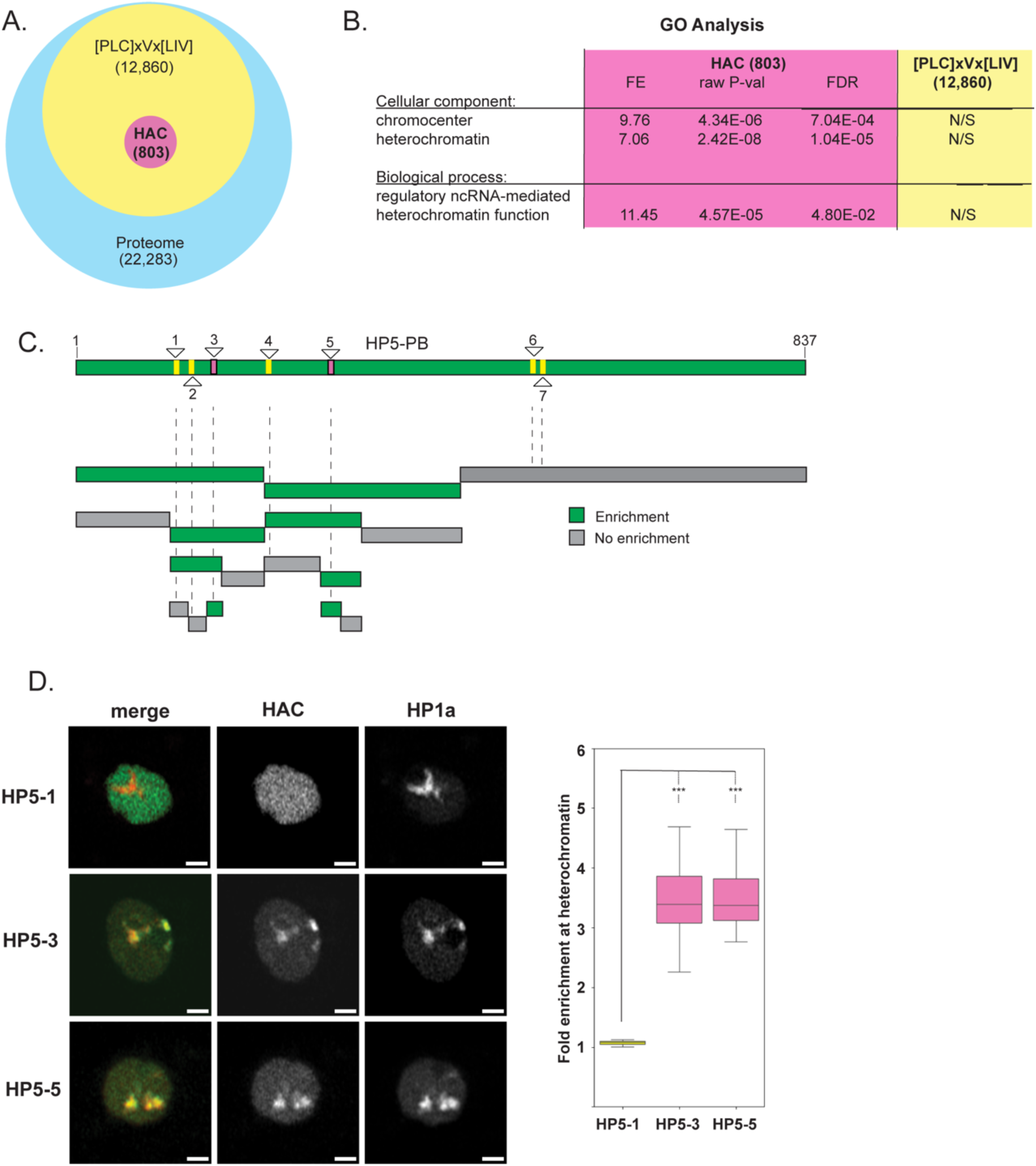
(A) Venn diagram showing the number of proteins harboring a PxVxL motif as defined by the indicated molecular signature versus those with the updated HAC signature. Compare with Supplementary Figure 1B. (B) GO analysis results showing the top annotations of “heterochromatin” and “chromocenter” among proteins containing the updated HAC signature, but not among proteins harboring the degenerate PxVxL motif; FE = fold enrichment above expected results, FDR = false discovery rate, N/S = not significant. (C) Diagram of HP5 isoform PB with locations of putative HP1a-binding sites based on either the PxVxL motif of hydrophobic-x-Val-x-hydrophobic (in pink) or the new HAC signature (in yellow) described in Figure 6B. Green rectangles indicate intact gene or fragments that show co-localization with HP1a in S2R+ cells after transient transfection; gray rectangles indicate lack of co-localization. (B) Representative images showing enrichment of HAC candidate peptides at HP1a-enriched heterochromatin (left). Quantification of HP1a co-enrichment of candidate HP5 HAC peptides (right). Scale bars = 2 microns.

To test the ability of the updated HAC signature to identify *bona fide* HP1a binding sites, we used a known HP1a interactor with no known binding sites, HP5, as a test case (**Figure 7C**). We identified two sequences in HP5 that matched our updated HAC signature (marked in pink), whereas 6 other sequences fit a more inclusive search comprised of a hydrophobic-x-valine-x- aliphatic signature (marked in yellow). HP5 was then subjected to a systematic fragment analysis to identify functional HP1a-binding regions in an unbiased manner, using imaging for HP1a co-localization as the readout (**Figure 7C**). Our results showed that only fragments containing the two binding sites predicted by our updated signature co-localized with heterochromatin **(Figure 7D),** suggesting these sequences are *bona fide* HP1a-binding sites. Imaging analysis of 25-mer peptides harboring these putative HACs displayed remarkably high enrichment at HP1a domains **(Figure 7D)**. Whereas individual mutations of the central residues of these two HACs (V159E or V294E) did not impact association of full-length HP5-PA with heterochromatin **(Supplementary Figure 7)**, a double mutation (V159E and V294E) dramatically reduced HP5 expression and abrogated all association with heterochromatin (see longer exposure). Therefore, our results indicate that HP5 is completely dependent on the predicted HAC motifs for associating with the HP1a condensate (**Supplementary Figure 7**). In conclusion, we propose that this updated signature, when combined with disorder prediction and conservation analysis, will be invaluable for identifying other HP1a interactors and regulators of the heterochromatin biocondensate.

## DISCUSSION

The HAC motif provides a widespread and evolutionarily conserved mechanism for HP1a binding and targeting of the heterochromatin domain. The results of this investigation expand our understanding of this HP1a-binding motif by exposing (1) the range of degeneracy of the core hydrophobic sequences that insert into the CSD dimer cleft, (2) the requirement of basic residues to stabilize HP1a association in Drosophila, and (3) the molecular interactions that modulate HAC affinity with the HP1a CSD and C-terminal tail. This led to expansion and modification of the original pentameric PxVxL motif into a larger consensus region **(Figure 6D)** consisting of 3-6 positively charged residues that interlace and surround a core of three alternating hydrophobic residues containing a central valine or isoleucine. This 9-mer basic consensus pattern is further flanked by a moderate frequency of both hydrophobic and positively charged residues, as well as low frequency of negatively charged and aromatic residues, over a 15-mer span that, altogether, generate the molecular interface for binding to the HP1a CSD dimer and CTE.

The HAC signature was determined through positional analysis of 6 previously identified PxVxL sequences in *Drosophila melanogaster*, 10 more sequences identified from HP1a- binding proteins through this study, and 157 sequences acquired from alignments of HP1a partner homologs in 11 other Drosophila species. We note that though most of the HAC homologs were untested in the peptide imaging assay, the positional and sequence conservation between most HAC homologs were readily distinguished in alignments, suggesting that protein-specific molecular interactions between HAC and HP1a are preserved between species. For example, the Su(var)3-9 HAC is 100% identical over its 15-mer length among 12 species of Drosophila, and the HP2-1 HAC is distinctive for containing a highly conserved Asn at the +3 position (**Supplementary Figure 6A**). We also validated the HP1a co-enrichment of 4 HAC homologs that represent some of the more divergent sequences (**Supplementary Figure 6B**). Finally, the HAC-interacting residues of the HP1a CSDs and C-termini are preserved among all Drosophila species included in this study (**Supplementary Figure 8A**). For these reasons, we conclude that HAC homologs represent functionally conserved HP1a-binding domains that greatly inform the HAC consensus signature. We do note that the high similarities in amino acid sequences within each group of HAC homologs cause overrepresentation of specific residues in the consensus analysis. Therefore, a more refined signature could be achieved through the inclusion of more homologs from other Drosophila species or discovery of more HACs in other HP1a-binding proteins.

Another predominant feature of HACs is their location within or in proximity to intrinsically disordered regions of proteins. Disorder prediction was conducted using the widely used PONDR VL-XT algorithm that integrates three neural networks trained on long (40+ residues), N-terminal, and C-terminal disordered regions of X-ray crystallographic data (Li *et al*., 1999; Romero *et al*., 2001). Though we cannot rule out the possibility that some HACs are located on exposed surfaces of globular secondary structures, we hypothesize that HAC association with intrinsically disordered domains confers flexibility and accessibility (Holehouse and Kragelund, 2024) that may be important for targeting the CSD dimer. The AlphaFold-Multimer model for the non-functional HAC candidate ADD1-1, which is predicted to contain a highly ordered structure, fails to position the central hydrophobic core of ADD1-1 at the center of the HP1a CSD dimer (**Supplementary Figure 8B**), despite containing a canonical motif (PxVxL). The binding region exhibited lower prediction confidence (pLDDT < 50) than functional peptides that contain highly disordered regions (pLDDT scores > 90). These structural prediction results suggest that the flexibility of the peptide, attributed to its disordered regions, may be necessary for proper fitting within the HP1a CSD dimer. We also detect the conservation of disorder correlation among putative HACs of other Drosophila species. HACs may therefore represent short linear motifs (SLiMs) or molecular recognition features (MoRFs) within disordered regions that mediate diverse functions (Hsu *et al*., 2013). Unlike well-defined globular structures, SLiMs and MoRFs possess weaker conservation due to the higher mutation rates found in disordered regions. Nevertheless, we identified positional biases in the surrounding residues of the central hydrophobic core of HACs that contribute to HAC binding with HP1a (**Figure 6D**). Aside from the functional role of flanking basic residues in modulating interactional strength with HP1a, we also observed a strong bias against acidic and aromatic residues when basic ones are absent, suggesting that their presence at these positions interfere with HAC function. Additionally, the location of HACs at highly mutable disordered domains and at paralogous locations in some protein homologs indicates that HACs may undergo relatively rapid evolutionary selection but become stably inherited once established. Our findings highlight the functional importance of intrinsically disordered domains as sites for protein targeting and the evolution of HACs.

Our findings also expand the molecular signature of the original PxVxL motif that forms the hydrophobic core of functional HACs. Though a trio of alternating hydrophobic residues is the classical feature of the PxVxL motif, there is currently confusion regarding the types of hydrophobic residues that can occupy each position in the motif, and whether this pentamer is sufficient for HP1a binding (Lechner *et al*., 2005; Meyer-Nava *et al*., 2020). Our search for the HAC molecular signature uncovers the range of amino acids that occupy each position of the hydrophobic core residues. Expectedly, we identified mostly valines at the central position of HACs, extending the previous structural prediction that only valine can be accommodated by the hydrophobic pocket of the Drosophila HP1a CSD dimer (Thiru *et al*., 2004). However, we also identified isoleucines at the central positions of Su(var)3-9 and Lhr HACs of Drosophila melanogaster, as well as in HACs of other homologs. Mutation of the central valine residue to glutamate or alanine has previously been shown to eliminate HP1a binding, and we demonstrate that mutation of a central isoleucine of a HAC similarly abrogates heterochromatin targeting (**Supplementary Figure 1C**). Meanwhile, we identify that the-2 position utilizes a much wider variety of hydrophobic residues, including polar cysteines, bulky phenylalanines, and conformationally rigid prolines. Conservation analysis of HACs across Drosophila evolution suggests that only the smallest and largest sidechain volumes of hydrophobic residues are excluded at the-2 position, namely glycine, tyrosine, and tryptophan. Additionally, our MD simulation shows that the sulfur in the cysteine sidechain at this position provides an additional hydrogen bond that may enhance HP1a binding. In contrast, we identify only small hydrophobic residues such as leucine, valine, and isoleucine to occupy the +2 position, though alignment analysis also identifies rare homologs containing glycine and methionine instead. This suggests that the +2 position is more sterically restrictive to larger hydrophobic residues compared to the-2 position. Altogether, this trio of residues comprise an asymmetric hydrophobic plug that is essential to recognize the conformation of the CSD dimer pocket, but which we also show is insufficient to target heterochromatin effectively *in vivo*. Our HAC fragment analysis (**Figure 3**) not only showed that the hydrophobic pentamer is insufficient for heterochromatin targeting, but also that the flanking 5 residues on each side of the pentamer are crucial for stable HAC association with heterochromatin.

Finally, we determined that electrostatic interactions are required for effective HAC-HP1a dimer binding, in addition to hydrophobic interactions. Despite clear exceptions, it is remarkable that there is a clear correlation between predicted Coulombic energy and measured fold-enrichment (**Figure 4D**), given the higher complexity of *in vivo* conditions and potential contributions from other factors not included in the simulations. In fact, the exceptions are informative about potential complexities in HP1-partner interactions. For example, Lhr-2 has larger attractive Coulomb energy (**Figure 4D**) but also greater hydrophilic surface areas than higher-ranked HP2-1 and HP4-3 peptides (**Figure 4B, Supplementary Figure 4D**). This suggests Lhr-2 contains a less stable hydrophobic pocket that could weaken HP1a interactions, despite its high electrostatic energy. Meanwhile, other exceptions, such as HP2-1, HP4-3, and Kdm4A, show high fold enrichment values despite lower Coulomb energy compared to other HACs (**Figure 4D**). HP2-1 possesses L-2, V0, and I+2, the top three amino acids on the hydropathy index (Kyte and Doolittle, 1982), whereas HP4-3 exhibits high hydrophobicity at L+5, and Kdm4A displays significant hydrophobicity at the flanking region, specifically at L-4 and L+5 (**Supplementary Figure 8C**). These peptides likely engage in more hydrophobic-driven interactions that compensate for their lower electrostatic energy. We should also note that electrostatic and hydrophobic interactions can negatively affect each other and be competitive (Dzubiella and Hansen, 2003, 2004; Wang, Friesner and Berne, 2010). For example, the Dmt HAC had weaker attractive Coulomb energy at the +1 and +4 positions than XNP and Piwi, likely due to the presence of isoleucine, which has the highest hydropathy index (Kyte and Doolittle, 1982), at the adjacent +2 and +5 positions (**Supplementary Figure 8D**). Electrostatic interactions from the +1 and +4 positively charged amino acids are key contributors to the HAC and CSD interaction (**Figure 5**). These results suggest that HAC peptides and CSD dimer interactions are mediated by both hydrophobic and electrostatic interactions, and that maintaining a moderate balance between hydrophobicity and charge is crucial to prevent competition between these types of interactions. Therefore, the contributions of electrostatic and hydrophobic interactions vary between HACs, and such structural flexibility could provide the variations in HP1a affinity that are important for each heterochromatin protein function.

Conservation analysis and MD simulations both indicate that positions-4,-3,-1, +1, +3, and +4 contain a strong bias for basic residues (lysine or arginine) that make electrostatic interactions with the HP1a CSD and C-terminal tail. In particular, we showed that the positively charged amino acids ±4 form a salt bridge interaction with Asp157 (**Figure 5A, B**). The positively charged amino acid at +1 plays an important role in maintaining the electrostatic force field by forming an H-bond with Arg197 of the CSD dimer and enabling the +3 positively charged amino acid to form salt bridges with Asp203 and Glu205 of the CSD (**Figure 5C, D,** and **E**). Additionally, we identified compensatory mechanisms when specific electrostatic interactions are absent at a given locus (+1, +3) (**Figure 5D, F**). The high conservation of positive charges in Drosophila HACs is made possible by the absolute conservation of interacting residues in Drosophila HP1a homologs, namely Asp157 of the CSD and Asp203 and Glu205 of the C-terminal tail (**Supplementary Figure 8A**). Acidic residues at positions 157 and 203 are also observed in HP1a homologs of humans, frogs, and fish, suggesting that HACs of these organisms also utilize basic residues for targeting heterochromatin (**Supplementary Figure 8E**). We note that, though HP1 C-terminal tails between these homologs are highly variable in length and sequence, they nevertheless are highly enriched for acidic residues that may partner with different positions of HACs for electrostatic interactions. We therefore propose that the inclusion of 3-6 basic residues within HAC positions-4 to +4 is an integral part of the HAC molecular signature in Drosophila HP1a, which may vary in content and position in non-Drosophila HACs.

CSD homodimer and the ADD1-1 are colored cyan and yellow, respectively. Residues at the - 2, 0, +2, +3, +4, +5, +6, +7, and +8 positions (in licorice drawing style) are colored orange, with the amino acid code indicated. (C) Visualization of surface hydrophilicity and hydrophobicity of XNP, H2-1, HP4-3, and Kdm4A in the side view were calculated using the Discovery Studio software. The hydrophilic to hydrophobic properties are colored from blue to brown. Each peptide is colored yellow. (D) Coulomb energy ± SD [kJ/mol] among the CSD dimer and-7 to +7 residues of XNP, Dmt, and Piwi. Orange circle: XNP, Green diamond: Dmt, and Blue square: Piwi. Electrostatic sequences at +1 and +4 are highlighted in orange; hydrophobic sequences at +2 and +5 are highlighted in pink. (E) Alignment of CSD and C-terminal tails of *D. melanogaster* HP1a with homologs from other species, including Homo sapiens, Xenopus, zebrafish, worm, and yeast, using a similar format as shown in (A).

Variations in the number and positions of basic residues in different HACs indicate that they act to modulate HP1a interactional strength. Though it is possible that there are HACs with fewer basic residues, we predict that these will exhibit a much lower affinity for HP1a and cannot be detected by our live imaging assay. Identification of an even more refined molecular signature may also be achieved with a wider conservation analysis beyond the 12 Drosophila species used here and more MD analysis for other types of peptides. To closely examine the molecular interactions of a HAC, we used the highest-enriching sequence of XNP to be a model peptide, by combining visualizations of interactions for residues from-7 to +7 with statistical observations of the MD simulations (**Figure 5I and Supplementary Figure 5I**). The last frame was initially used to make the interaction map, which was then adjusted by comparing it with the statistical results, such as Coulomb energy, to obtain a more accurate interpretation. Based on the interaction map, conservation analysis, and Coulomb energy, we can infer that the HAC positions between-4 and +4 form the strongest molecular interactions, extending the previously recognized 5-mer signature of PxVxL. As shown in **Figure 5I**, each amino acid has distinct roles in the interactions between the peptide and the CSD dimer+CTE, and these roles can change depending on the neighboring sequences. We note, for example, that homologs of each HAC tend to possess additional conserved molecular features within their group that are beyond the HAC consensus sequence (**Supplementary Figure 6A**), pointing to protein-specific HP1a interfaces. A more comprehensive mutational study of such protein-specific residues could uncover additional mechanisms of peptide interaction. Alternatively, factors other than hydrophobicity and electrostatic forces could also be contributing to HAC-HP1a interactions, such as post-translational modifications of peptides or HP1a.

But why would an affinity hierarchy exist for protein interactions with HP1a? HAC structures likely reflect the specific requirements of heterochromatin proteins to achieve different levels of enrichment or mixing in heterochromatin. For example, rare proteins may contain high-affinity HACs for efficient targeting, whereas more abundant proteins, or those containing additional domains that interact with other heterochromatin factors, will not require such high affinity. Alternatively, structurally important heterochromatin proteins likely contain HACs that stably associate with HP1a, whereas enzymes or proteins that only interface transiently with heterochromatin may require lower affinities. For example, HP5 with its two strong HACs could form highly stable heterochromatin structures or even crosslink two HP1a dimers, whereas the Su(var)3-9 methyltransferase likely requires only transient association with HP1a as its chromodomain is expected to bind H3K9me2/3 in heterochromatin (Müller *et al*., 2016). HAC affinities could therefore facilitate how HP1a organizes heterochromatin architecture by directing the activity of multiple complexes to execute diverse functions in this complex, dynamic biocondensate.

A hallmark of biocondensates is the accumulation of multivalent interactions between their constituents to drive partitioning from the surrounding environment (Banani *et al*., 2017). In heterochromatin, this is achieved by the relatively abundant HP1 protein that centrally organizes the accumulation of other structural condensate proteins through its abilities to bind the heterochromatin-specific histone mark H3K9me2/3, homodimerize, and recognize HACs (Janssen, Colmenares and Karpen, 2018). This inherent demixing ability also effectively diminishes, if not outright excludes, other nuclear proteins from the condensate (Banani *et al*., 2017). Nevertheless, the underlying heterochromatic DNA polymer is not precluded from pan-nuclear processes such as transcription, replication, and repair. Therefore, heterochromatin must either (1) employ dedicated factors to carry out such functions, (2) recruit factors from such processes to access the interior, (3) transiently or locally dismantle the condensate for said processes to proceed, or 4) a combination of these mechanisms. HACs can provide the means to carry out at least the first two potential mechanisms, though controlled destabilization of heterochromatin may also utilize an activatable HP1a-interacting protein. The HAC-containing proteins SuUR and Mu2 appear to represent such factors that regulate DNA replication and repair in heterochromatin, respectively, as SuUR is known to suppress polytenization of heterochromatic domains (Belyaeva *et al*., 2006) and Mu2 is an early DNA damage marker that recruits downstream repair proteins (Dronamraju and Mason, 2009).

HACs contribute to the multivalency of heterochromatin by facilitating the mixing of HP1a with multiple proteins. Conversely, HP1a CSD dimerization is enhanced through binding HACs, as has been previously shown with HP2-1 and Piwi peptides (Mendez *et al*., 2011). Whether higher HP1a affinity correlates with enhanced dimerization is currently unknown, but HACs with different binding strengths, in combination with properties of linked domains in full length versions, could regulate heterochromatin condensate formation, dissolution, and/or material properties. Interestingly, mammalian HP1 phase separation properties were shown to be differentially regulated by 2 different PxVxL-containing peptides, further indicating that HAC-binding alters the biophysical properties of HP1 directly (Larson *et al*., 2017). Our results also indicate that HACs show enrichment throughout the heterochromatin domain, even though the full-length proteins encoding certain HACs, such as SuUR and XNP, normally display smaller subdomains of enrichment within heterochromatin (Swenson *et al*., 2016). Subdomain formation within heterochromatin must therefore be mediated by the concerted action of HACs and other types of interaction domains present in full length proteins. One possible mechanism is that such proteins utilize HACs to gain access to the heterochromatin interior and then, through strong self-interacting domains or protein partners, form subdomain condensates within heterochromatin. Through a similar pairing of self-interacting domains and particularly weak affinity HACs, other nuclear bodies, such as those that facilitate transcription, replication, and repair, may also gain access to the heterochromatin periphery that, upon activation, seed the formation of complexes to carry out functions (Rajshekar *et al*., 2023).

We foresee that our improvements in identifying HAC molecular signatures will help identify such ‘seed’ components for various nuclear processes, as well as focus attention on intrinsically disordered regions of various proteins to contain access codes to different biocondensates in the nucleus (King *et al*., 2024). Whether all condensates contain short linear motifs to facilitate the targeting of their components is not known, but our experimental and computational approaches could provide a framework to identify molecular grammar utilized by other types of biocondensates. Based on our findings with HACs, we propose that affinity hierarchies by short disordered motifs may comprise a general mechanism by which various biocondensates coordinate structural and transient components, as well as access shared nuclear components and even interface with other condensates.

## ONLINE METHODS

### Plasmid Construction

Oligonucleotides of putative HAC sequences (IDT) were cloned into a pCOPIA vector containing a N-terminal mNeonGreen fluorescent tag and a myc nuclear localization signal. HACs used for testing variable lengths of flanking sequence contained GGGSG in place of sequences-12 to-8 and lacked positions +8 to +12, or GGGSGGSGGG in place of-12 to-3 and lacked positions +3 to +12. Corresponding control constructs contained GGGSG or GGGSGGSGGG after the mNeonGreen tag and myc NLS. Full-length genes for HP1a, XNP, SuUR, Lhr, Su(var)3-9, HP4, Mu2, and HP5, along with HP5 gene fragments, were cloned into pCOPIA vectors containing N-terminal mCherry, mScarlet-1, mCitrine, or mGFP fluorescent tags as described previously (Colmenares *et al*., 2017). Mutants for HACs and full-length genes were generated using a one-step directed mutagenesis protocol (Liu and Naismith, 2008).

### Bioinformatics

25-amino acid sequences containing a central pattern of hydrophobic-x-Val/Ile-x-hydrophobic consensus were identified as putative HACs using custom Perl scripts. Hydrophobic residues included Ala, Cys, Phe, Ile, Leu, Met, Pro, Val, Trp, and Tyr, whereas x = any amino acid. Homologs of *Drosophila melanogaster* protein sequences were obtained using standard NCBI protein-protein BLAST by retrieval of sequences with the highest homology scores for each of the other 11 species of Drosophila (taxid:7215), using the non-redundant protein database. Conservation of putative HACs was determined by testing whether the same consensus pattern is detected to align with the putative *Drosophila melanogaster* HAC sequence after multiple sequence alignment of full-length homologous proteins with Clustal Omega (Madeira *et al*., 2024). Initially, only sequences containing a hydrophobic-x-Val/Ile-x-hydrophobic pattern that spatially aligned with the putative Drosophila melanogaster HAC, as determined by Clustal Omega with default settings, were scored positively to be conserved (see Supplementary Figure 2) and are shown on Figure 2. However, re-analysis of HP1a-binding protein homologs for the final HAC consensus motif (see below) revealed additional putative HAC sequences in other parts of SuUR, Mu2, and HP2 homologs that do not align with the Drosophila melanogaster HAC and may represent paralogous HACs. These additional sequences were included in the analysis for Figure 6 and Supplementary Figure 6. Disorder prediction was analyzed using PONDR-VLXT (Romero *et al*., 2001). Positional frequency of each amino acid was analyzed among groups of HACs using custom Perl scripts. The final HAC consensus used for motif search from positions-7 to +7 are as follows: [˄WF] [˄DEW] [˄WY] [˄STFWY] [˄DEFW] [FPLICVMA] [˄DEFWY] [VI] [˄DEFWY] [LIVGM] [˄WY] [ KRLN] [˄DEW] [ ˄FWY], where brackets contain all possible amino acids at each position, or all the excluded amino acids if marked with “˄.”

### Tissue culture manipulation and image acquisition

S2R+ embryonic tissue culture cells were maintained in Schneider’s medium (Thermo) with 10% fetal bovine serum (Gemini) and 1% antibiotic-antimycotic (Gibco) at 25°C. Transient co-transfections with mNeonGreen-tagged constructs and mScarlet-I-tagged HP1a were conducted using the TransIT-2020 reagent (Mirus). Cells were mounted 72 hours later on chambered coverslip slides (Ibidi) and live imaging was performed using a Zeiss LSM880 Airyscan microscope with 63X oil immersion objective at room temperature. A mNeonGreen tag containing only a myc NLS was co-transfected with mScarlet-I-HP1a as a negative control.

Transient co-transfections of fluorescently-tagged full-length HP1a-interacting proteins with HP1a were similarly conducted and images were collected using an Applied Precision Deltavision microscope or a Zeiss LSM880 Airyscan microscope with 63X oil immersion objective. All results are based from at least two biological replicates.

### Image analysis

Raw images from the Zeiss Airyscan were processed using ZEN black edition software, whereas Deltavision images were processed using SoftWoRx. All representative manuscript images were formatted using Fiji software. To calculate heterochromatin enrichment, 3-D Airyscan images of HP1a were segmented into nucleus, chromocenter, and nucleolus volume masks by manually setting the intensity thresholds in Arivis Vision4D software. A low intensity threshold setting selected the nucleus, a high setting was used to segment the chromocenter, while the negative space lacking HP1a signal adjacent to the chromocenter was isolated by the subtraction of HP1a nuclear masks with holes from those without holes. The nucleolar mask was dilated 3-4 pixels to avoid potentially measuring nucleolar intensity signal of mNeonGreen, which sometimes accumulated in the nucleolus in certain constructs, including the negative control. The mean chromocenter signal intensity of mNeonGreen was therefore measured only in chromocenter regions that did not overlap the dilated nucleolar volume, and its chromocenter enrichment was calculated by dividing the mean chromocenter intensity with the mean nuclear intensity. 10-36 nuclei were measured for each construct tested from 2 biological replicates. Student’s t test, assuming two-sample tails and unequal variance, was calculated.

### Alphafold Multimer-predicted computational model

AlphaFold-Multimer (Evans *et al*., 2022) was employed to develop the computational models of the HAC peptide and HP1a CSD dimer with C-terminal tails. The sequences for the Drosophila HP1a CSD dimer with C-terminal tails and flanking residues, sourced from the protein database (PDB ID: 3P7J) (Mendez *et al*., 2011), served as inputs for AlphaFold-Multimer computational model predictions: MKKHHHHHHEQDTIPVSGSTGFDRGLEAEKILGASDNNGRLTFLIQFKGVDQAEMVPSSVANEKIPRMVIHFYEERLSWYSDNEDKK. The 16 heterochromatin-enriched peptides identified by our experiments (**Figure 1E**) were selected for computational modeling to investigate the key features of the HP1a-peptide binding. The 25-amino-acid sequences for all 16 peptides are shown in Supplementary Method Table 1. The “pdb100” was used for the template mode. The model type was set to “alphafold2_mmultimer_v3”. The highest-ranked model for each peptide, based on pLDDT (predicted Local Distance Difference Test) scores, was used for the following computational analysis. All predicted models, except for Odj-1, showed high pLDDT (predicted Local Distance Difference Test, scaled 0-100) and pTM (predicted Template Modelling, scaled 0-1) scores, representing confidence in the local and overall structure of the complex, respectively (**Supplementary Method Table 1**). The 15 out of 16 peptide models positioned their central core region at the center of the CSD dimer (**Figure 3A, Supplementary Figure 3A-O**). While the disordered edge regions of the complex showed low prediction accuracy, the key binding sites between the peptide and CSD dimer had very high confidence with over 90 pLDDT scores (**Supplementary Figure 3A-O**). Odj-1 was the only model that mispositioned the peptide and had low prediction accuracy even in the binding sites with 60-80 pLDDT scores. Due to its low prediction confidence, the Odj-1 model was excluded from further computational analysis. The 15 peptide models, except for Odj-1, were used for the following molecular dynamics simulation analysis.

### Molecular dynamics simulation

GROMACS 2020.4 software package (Pronk *et al*., 2013; Abraham *et al*., 2015; Páll *et al*., 2015; Lindahl *et al*., 2020) was used for the all-atom molecular dynamics (MD) simulation with the CHARMM36 force field (Best *et al*., 2012; Huang and Alexander D MacKerell, 2013). The developed model of the HAC peptide and HP1a CSD dimer with C-terminal tails was inserted into a cube box (13*13*13 nm^3^) with periodic boundary conditions applied in all three directions. The CHARMM-modified TIP3P water model was employed to solvate the system (Boonstra, Onck and van der Giessen, 2016). The sodium and chloride counter ions were added to the system to obtain the 150 mM concentration of NaCl, a condition where chromatin is known to be aggregated (Huang and Cole, 1984; Shakya *et al*., 2020). Energy minimization was conducted for 50,000 steps with an energy tolerance of 1000 kJ⋅mol^-1^nm^-1^ to eliminate no steric clashes and ensure proper geometry of the system. The target temperature and equilibration of the solvent and ions in the system were achieved by employing NVT (isothermal-isochoric) and NPT (isothermal-isobaric) ensembles, each for 5 ns with a time step of 1 fs, with V-rescale thermostat at a constant temperature of 298 K. The Parrinello-Rahman barostat was used during NPT ensembles to maintain a pressure of 1 atm. The damping parameters were set to 0.1 ps for the V-rescale thermostat and 2 ps for the Parrinello-Rahman barostat. The heavy atom’s positions of the protein were constrained during NVT and NPT equilibration to prevent the additional variable of structural changes in the protein. Subsequently, protein relaxation was employed for 250 ns (with 2 fs time steps) as the product step without any position restraints in the equilibrated system. The MD simulations were performed using NVIDIA V100 SMX2 GPU at the San Diego Supercomputer Center. The particle mesh Ewald method was used for long-range electrostatic interactions. The 1.0 nm cutoff was used for short-range electrostatic and van der Waals interactions.

### Analysis of Molecular Dynamics Simulations

The trajectory files saved every 10 ps were used for the analyses. The HP1a CSD dimer (amino acids 141-201) in Piwi and XNP peptide models was stable throughout the 250 ns simulations (**Supplementary Method Figure 1C**). The HAC peptides of Piwi and XNP from-7 to +7 amino acids reached a stable state after around 175 ns (**Supplementary Method Figure 1D**). Data from 200 ns to 250 ns after reaching the stable state was used for computational analysis. The simulation was visualized using VMD software (Humphrey, Dalke and Schulten, 1996). All analyzed results were plotted using Python.

Root Mean Square Deviation (RMSD): RMSD of the HAC central pentamer from-2 to +2 for the last 50 ns was calculated using “gmx rms” GROMACS tool built-in function.

Root Mean Square Fluctuation (RMSF): RMSF of the XNP and Piwi of 25-residue HAC sequences (-12 to +12) for the last 50 ns was calculated using “gmx rmsf” GROMACS tool built-in function. Averaged RMSF of the distal flanking 10 sequences (-12 to-8 and +8 to +12), nearby flanking 10 sequences (-7 to-3 and +3 to +7), and central 5 sequences (-2 to +2) for XNP and Piwi was calculated, respectively. Tukey test was employed to compare differences between multiple groups (p-values < 0.05).

Hydrophobic and charge distribution: The hydrophilic and hydrophobic distribution and electrical charge distribution of the HP1a-interacting XNP regions were visualized using BIOVIA Discovery Studio (Dassault Systèmes, USA) (*Discovery Studio*, 2023).

Hydrophilic surface area: The hydrophilic surface area of the HAC central pentamer (-2 to +2) and the pentamer with nearby flanking (-7 to +7) were computed for the last 50 ns of the MD simulations using “gmx sasa” GROMACS tool built-in function for oxygen and nitrogen atoms.

Coulomb energy: The short-range Coulomb energy between the HAC peptide and HP1a CSD dimer with C-terminal tails with a 1.0 nm cutoff distance was calculated for the last 50 ns of the MD simulations using “gmx energy” GROMACS tool built-in function. The Coulomb energy for each individual residue was calculated in the same manner. The frequencies of the coulombic energy with kernel density estimation were plotted using Python.

BIOVIA (*Discovery Studio*, 2023) visualization of hydrophilicity/hydrophobicity and surface charge distributions: The hydrophobicity on the peptide is derived from the Kyte-Doolittle scale. For the charge calculation, partial charges assigned to each surface atom are used. The algorithm details can be found in BIOVIO Discovery Studio (*Discovery Studio*, 2023).

VMD (Humphrey, Dalke and Schulten, 1996) visualization of hydrogen bonds (H-bonds) and salt bridge interactions: H-bonds were colored in green when the distance between the possible acceptor (OH and NH groups) and donor (O and N atoms) pair was within 0.35 nm, and the angle between the hydrogen, oxygen, and nitrogen was below 30°. Salt bridge interactions were colored in orange when the distance between the side-chain carbonyl oxygen of Asp or Glu and the side-chain nitrogen atoms of Arg, Lys, or His is within 0.4 nm.

2D molecular interaction map: The 2D molecular interactional map of XNP and HP1a CSD dimer with C-terminal tails was created using Adobe Illustrator (Adobe Inc., USA) (*Vector Graphics Software – Adobe Illustrator*, no date) based on the 2D diagrams for each residue analyzed by BIOVIA Discovery Studio (Dassault Systèmes, USA) (*Discovery Studio*, 2023).

## Acknowledgements

Many thanks to members of the Karpen lab and Dr. Grace Y.C. Lee for their valuable feedback.

We thank Dr. Ksenia V Krasileva and Dr. Daniil M. Prigozhint for their helpful discussion regarding AlphaFold-Multimer modeling, the Japan Student Services Organization (JASSO) Student Exchange Support Program (Graduate Scholarship for Degree Seeking Students), the SAITAMA GO GLOBAL STUDENT scholarship, and the Hearts To Humanity Eternal (H2H8) graduate research grant for supporting Shingo Tsukamoto’s doctoral studies. This research used resources of the Advanced Cyberinfrastructure Coordination Ecosystem: Services & Support (ACCESS) super-computing facilities, supported by the National Science Foundation (NSF) Grant No. BIO230220. The Karpen lab is supported by the National Institutes of Health (R35GM139653).

---

**Supplementary Figure 1.**
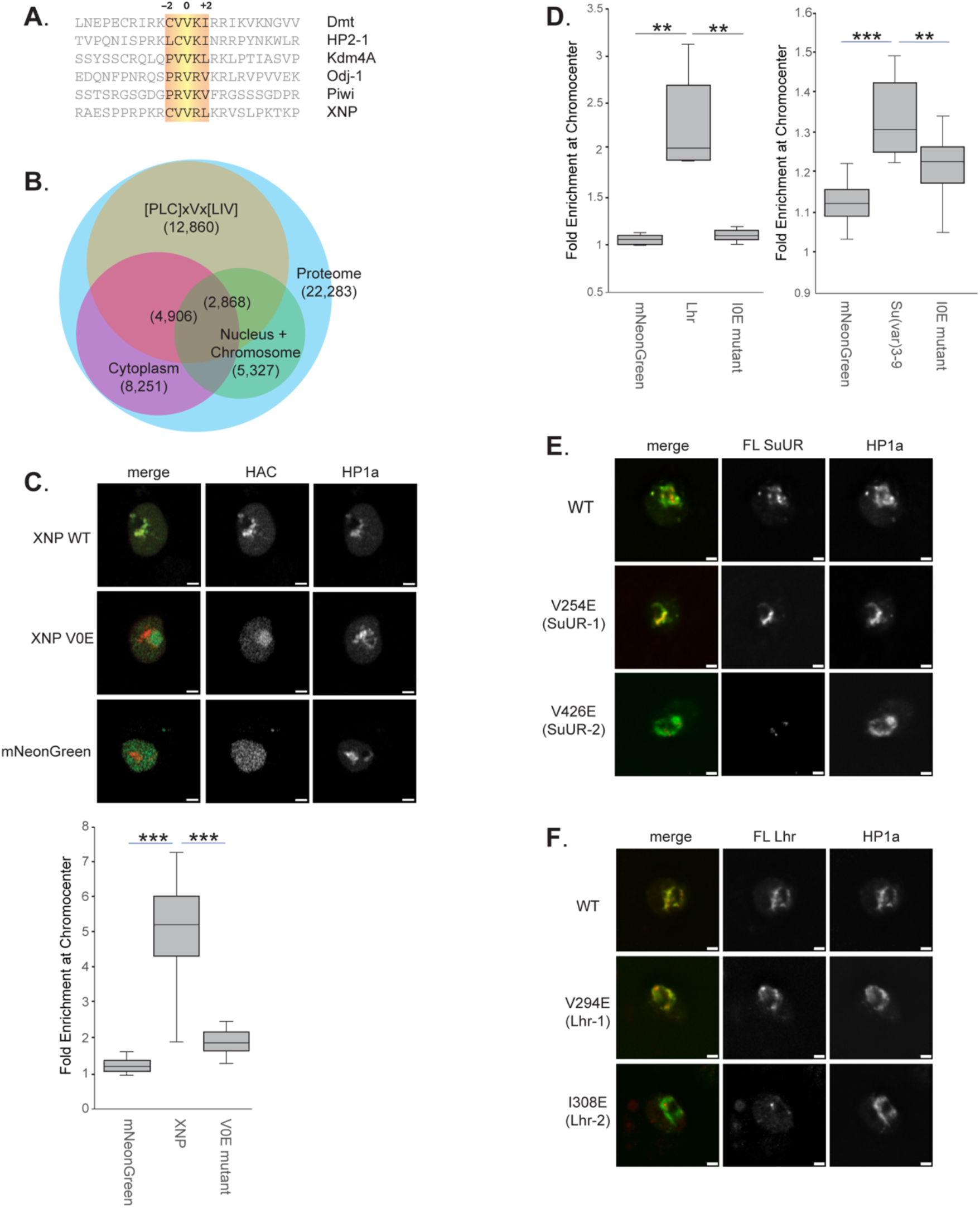
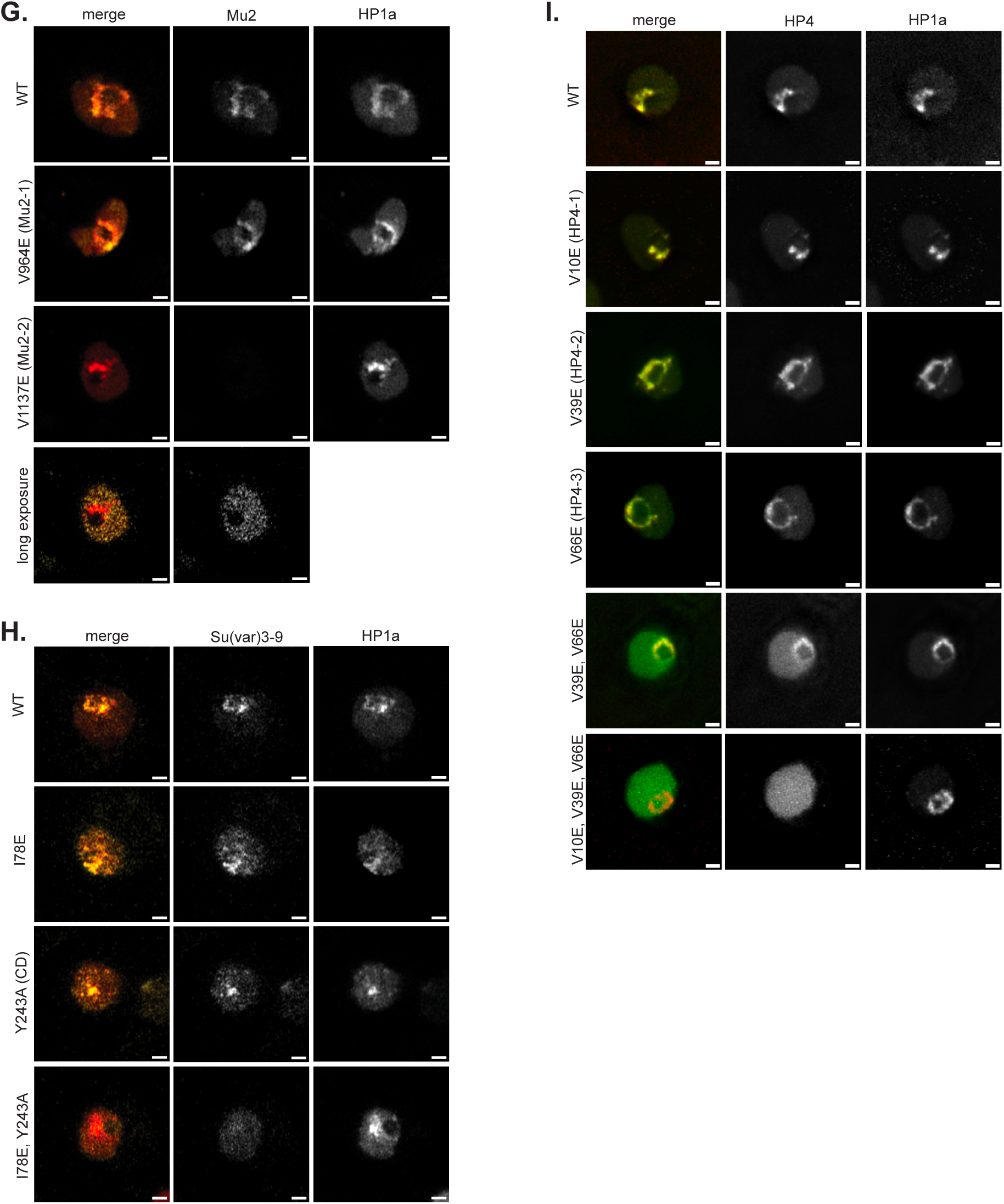
(A) Sequence alignment of 6 known PxVxL domains (highlighted in yellow) from published research with positions-2, 0, and +2 marked. (B) Venn diagram of proteins containing degenerate PxVxL consensus and their association with GO-annotations of nucleus/chromosome or cytoplasm components. Number of proteins within each group is shown as well as those containing the motif and annotated to be nuclear/chromosomal or cytoplasmic. (C) Representative images of mNeonGreen-tagged wildtype and mutant XNP HAC peptide in relation to mScarlet-I-tagged HP1a in S2R+ cells (top). Mutant XNP HAC has its central residue mutated from Val to Glu (V0E). Quantitation of fold-enrichment in HP1a-rich heterochromatin condensates by XNP HAC constructs and a control containing only the fluorescent tag (bottom). *** = p-value<0.001; ** = p-value<0.01. (D) Validation of putative Lhr and Su(var)3-9 HACs by mutation of central Ile to Glu (I0E). Representative images of full-length SuUR (E), full-length Lhr (F), and full-length Mu2 (G) with and without V0E mutations at putative HAC sites. A longer exposure for mutant Mu2 is included. (H) Representative images of full-length Su(var)3-9 with or without HAC and chromodomain (CD) mutations. (I) Representative images of full-length HP4 with or without individual or combinatorial HAC mutations. Scale bars = 1 um.

**Supplementary Figure 2:**
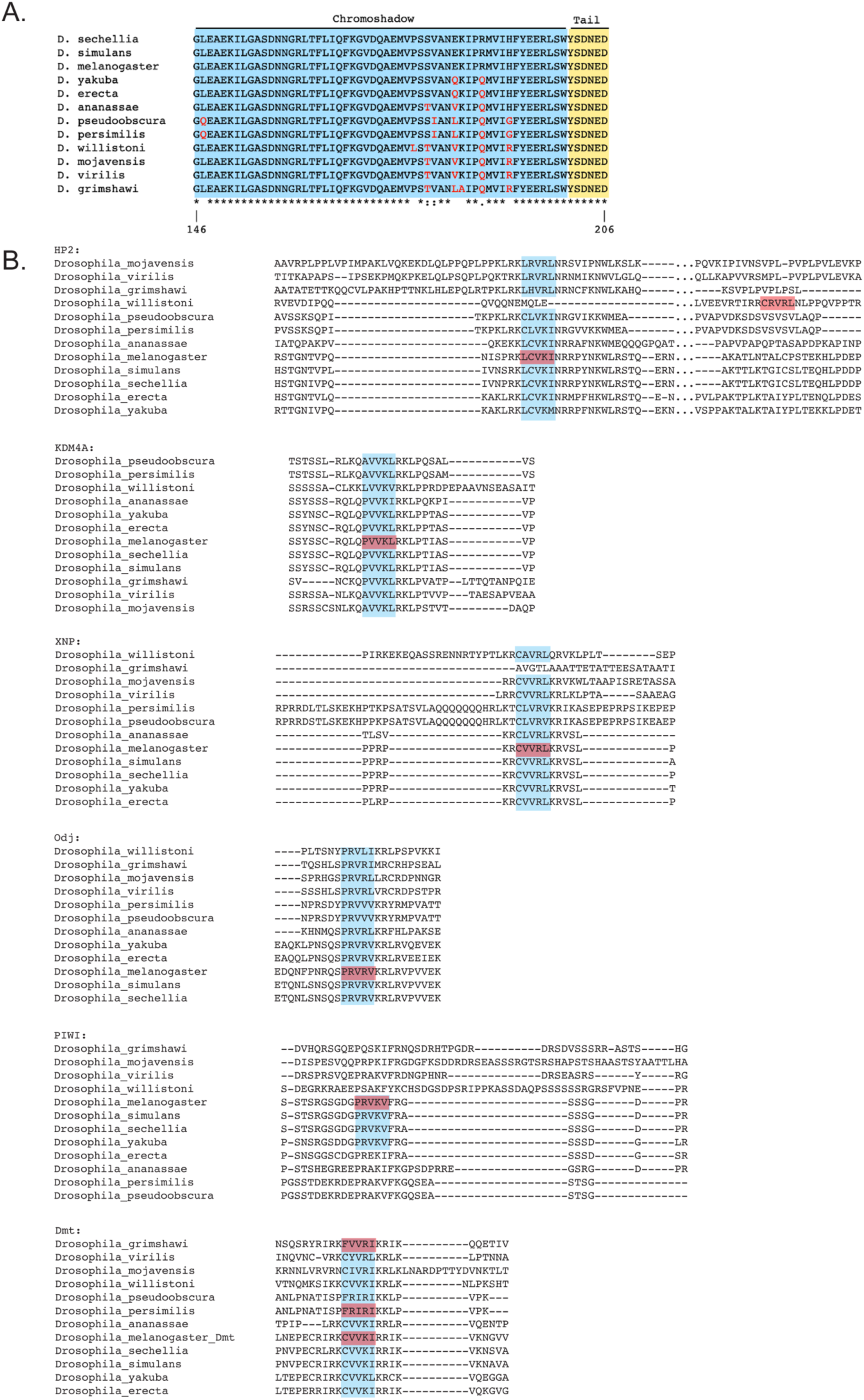
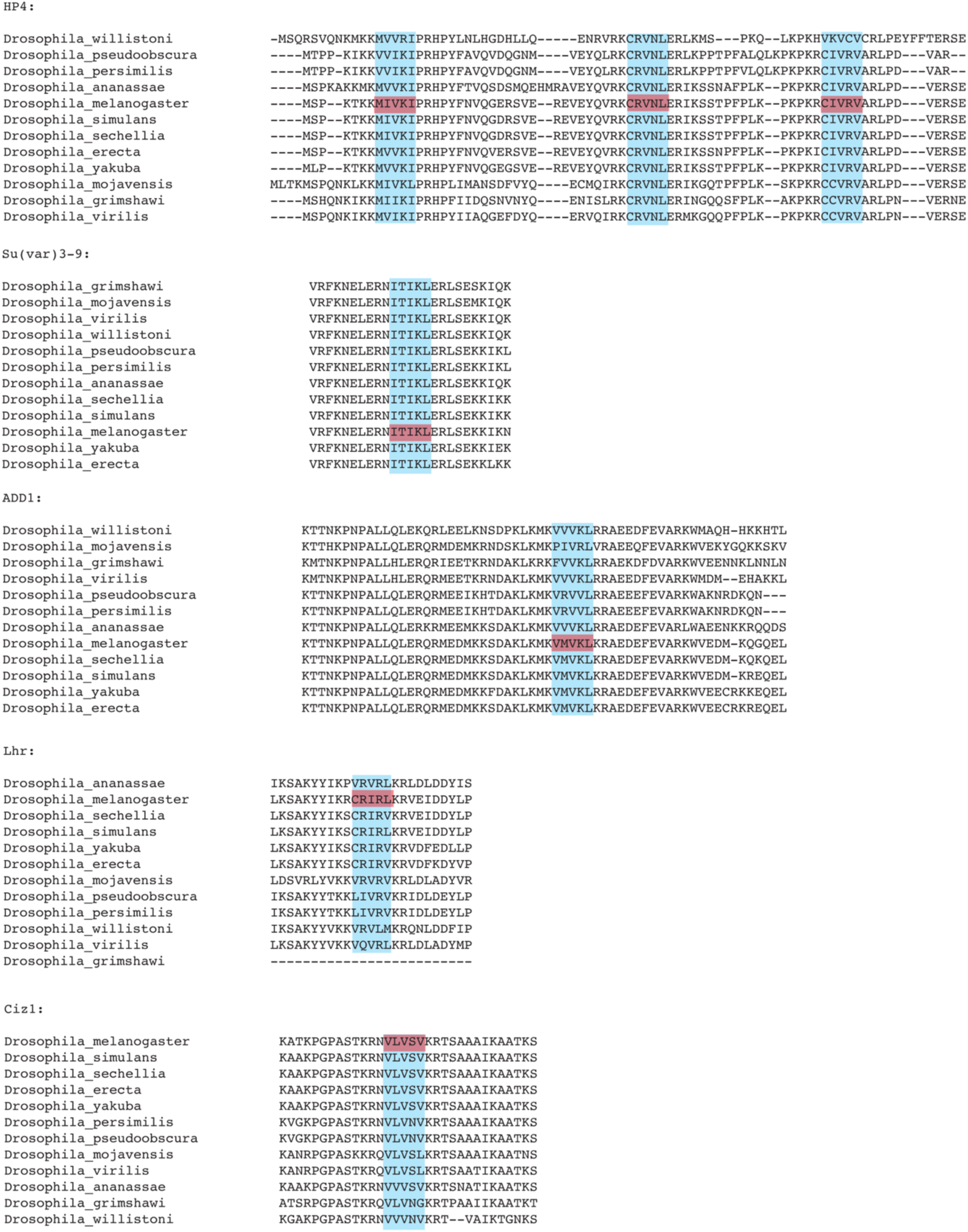
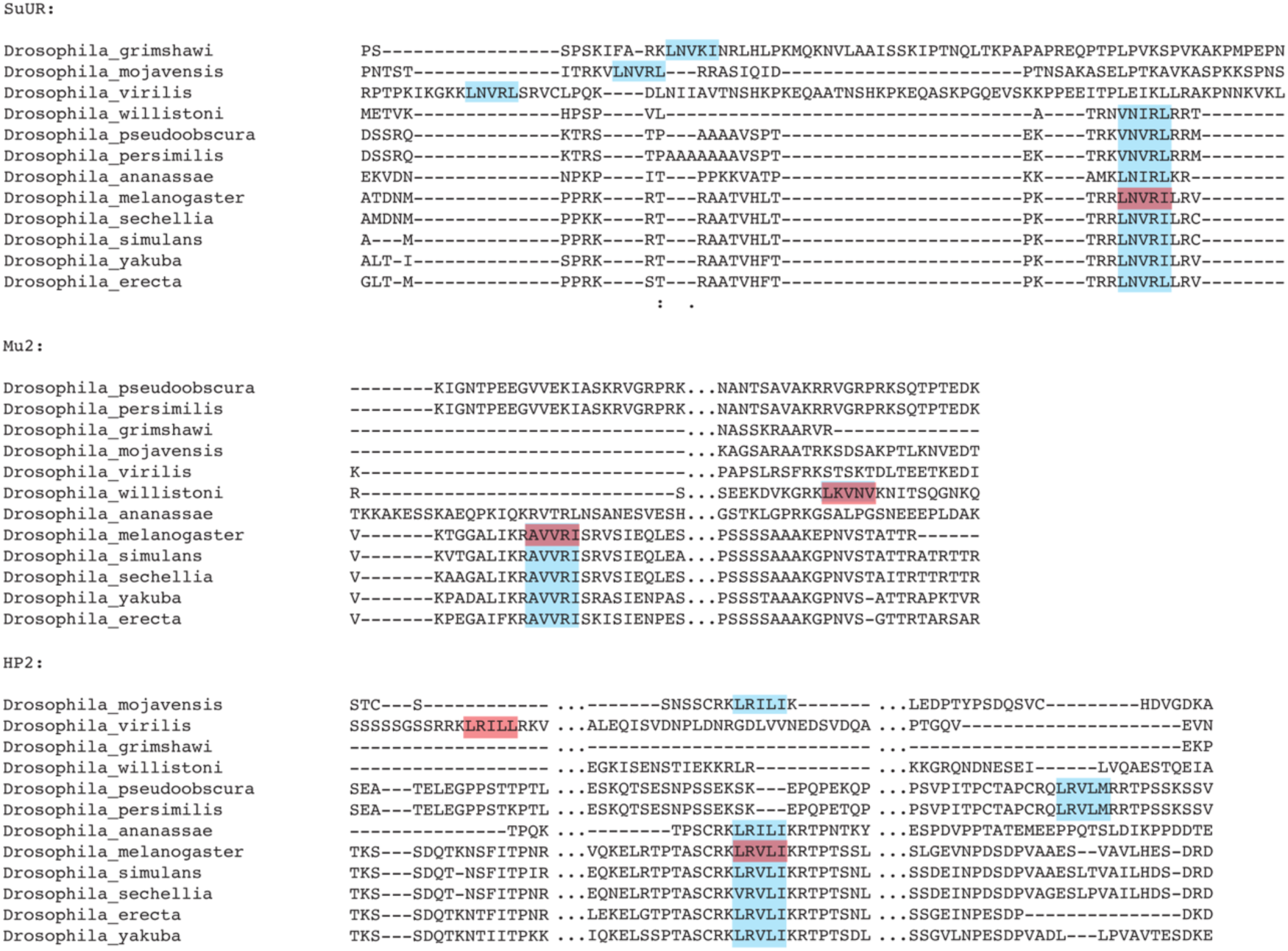
(A) Sequence alignment of HP1a CSD (highlighted in blue) and CTE (highlighted in yellow) for 12 Drosophila species. Divergent residues are marked in red. (B) Alignments of HP1a-binding proteins containing HACs across 12 Drosophila species. Pentameric core domains of HACs are highlighted in blue; HACs tested to accumulate in heterochromatin are highlighted in red. Some HAC sequences were manually curated as alignments of SuUR, Mu2, and HP2 homologs did not align with the Drosophila melanogaster HAC sequences and may represent paralogous HACs.

**Supplementary Figure 3:**
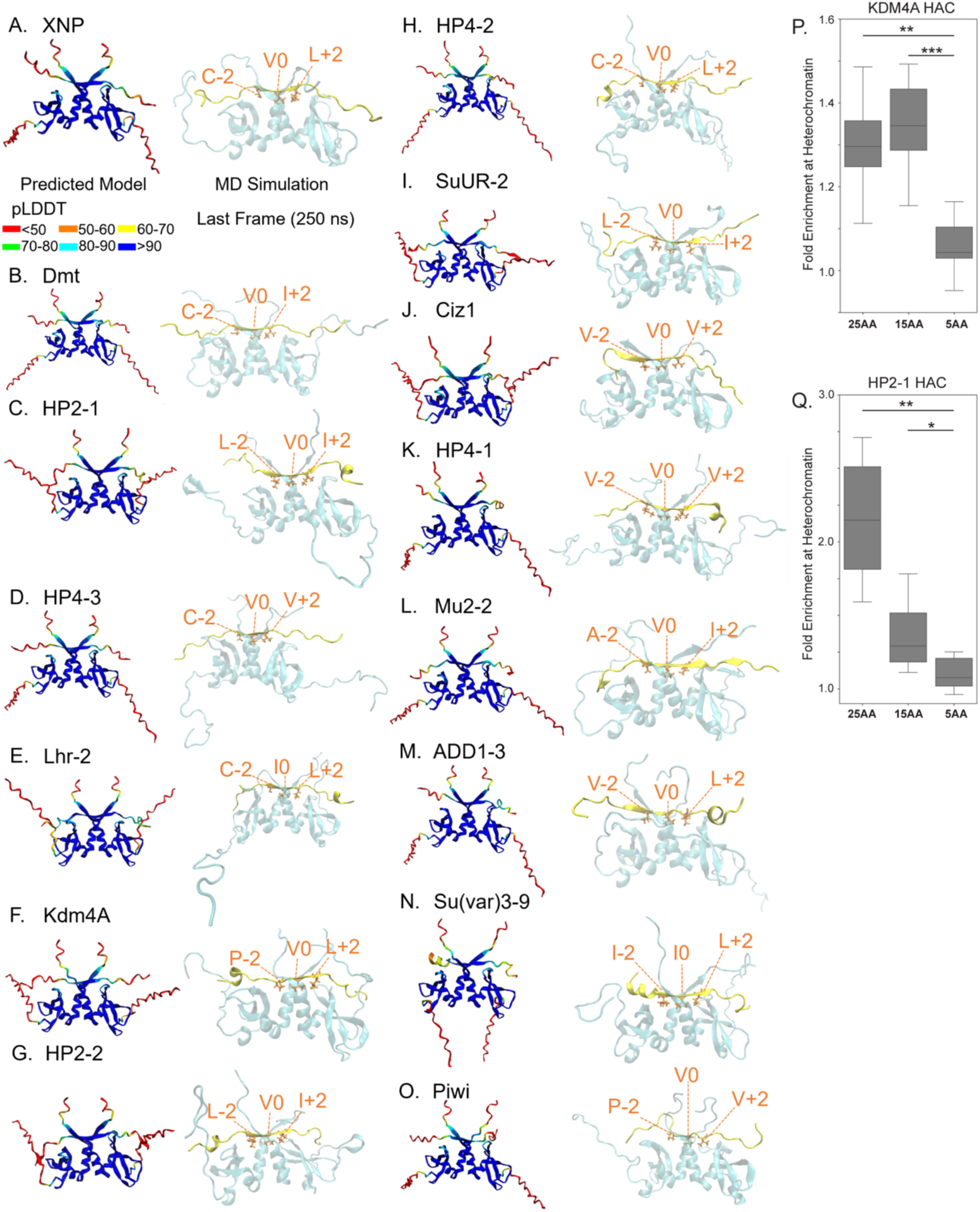
(A-O) The Alphafold Multimer-predicted computational model (left) and visualized MD simulation image of the last frame at 250 ns (right) for the CSD dimer and 15 peptides. The Alphafold Multimer-predicted structure is colored based on pLDDT (predicted local distance difference test) scores. Red: pLDDT<50, Orange: 50<pLDDT<60, Yellow: 60<pLDDT<70, Green: 70<pLDDT<80, Skyblue: 80<pLDDT<90, and Blue: 90<pLDDT. CSD homodimer and the peptide are colored cyan and yellow, respectively. Residues at the-2, 0, and +2 positions (in licorice drawing style) are colored orange, with the amino acid code indicated. (P-Q) Quantitation of HP1a co-enrichment at heterochromatin domains for the tested KDM4A and HP2-1 HAC constructs, respectively. 25AA = full-length native sequence containing core region surrounded by 10 residues on each flank; 15AA = core region surrounded by 5 residues on each side; 5AA = core region only. *** = p-value < 0.001; ** = p-value < 0.01; * = p-value < 0.05.

**Supplementary Figure 4:**
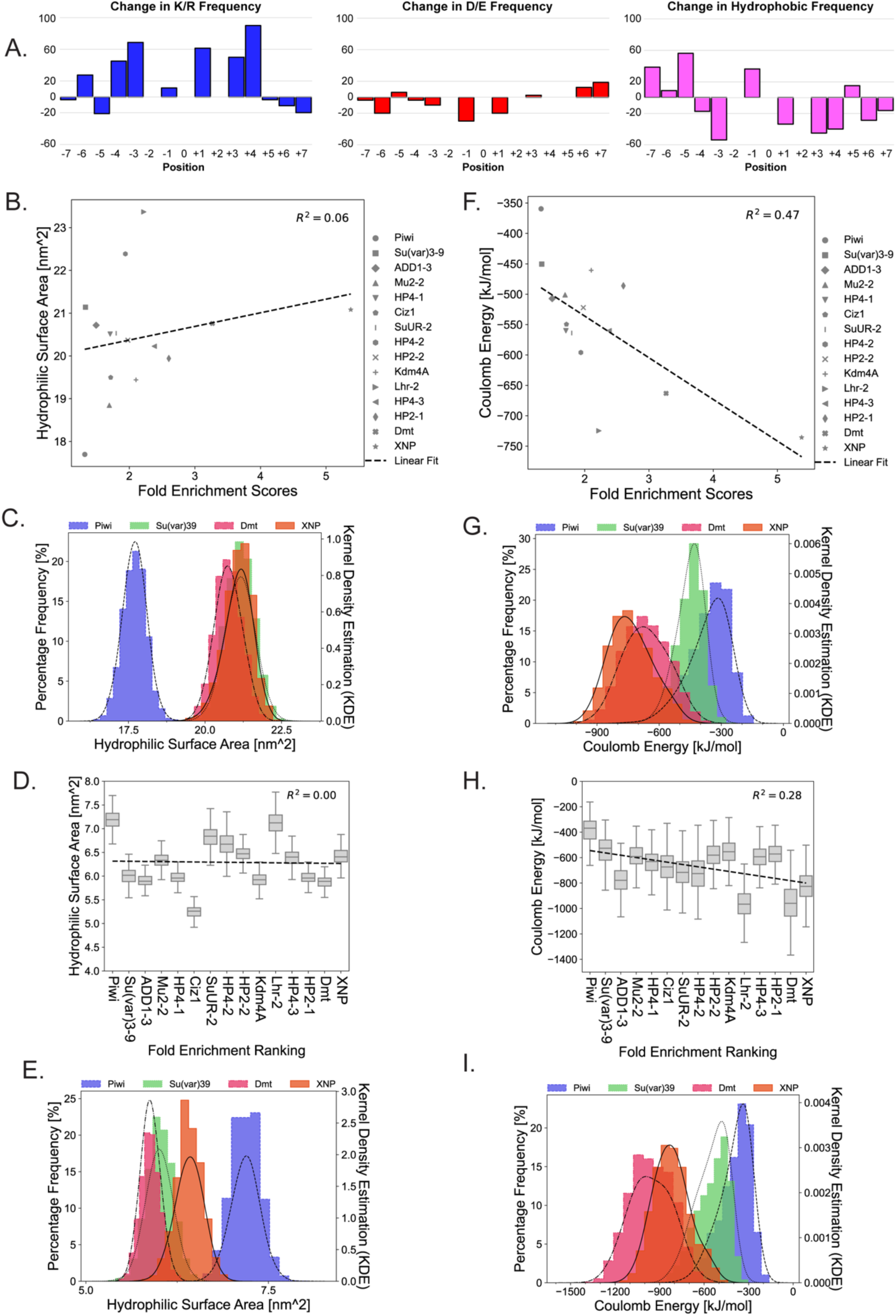
(A) Bar graphs showing differences in percent frequency of positively charged residues (left in blue), negatively charged residues (center in red) and hydrophobic residues (right in pink) between HAC peptides and non-HACs. (B) Hydrophilic surface area of sequences from-7 to +7 of 15 HACs between 200 to 250 ns of the MD simulations are plotted according to the fold HP1a co-enrichment values (Figure 1E). The R-squared value is shown. (C) Histogram showing the percentage frequency of hydrophilic surface area for sequences from-7 to +7 of HACs for Piwi, Su(var)3-9, Dmt, and XNP between 200 to 250 ns of the MD simulations. Piwi: Blue with a dash line. Su(var)3-9: Green with a dotted line. Dmt: Pink with a dash and dotted line. XNP: Orange with a solid line. Histogram was fitted by kernel density estimation (KDE). (D) Hydrophilic surface area ± SD [nm^2^] of sequences from-2 to +2 of 15 HACs between 200 to 250 ns of the MD simulations. The peptides are plotted according to ranking in terms of fold HP1a co-enrichment. The R-squared value is shown. (E) Histogram showing the percentage frequency of hydrophilic surface area for pentameric core sequences from-2 to +2 of HACs for Piwi, Su(var)3-9, Dmt, and XNP between 200 to 250 ns of the MD simulations, using the same format as C. (F) Coulomb energy of 15 HAC sequences from-4 to +4 between 200 to 250 ns of the MD simulations are plotted according to HP1a co-enrichment values. The R-squared value is shown. (G) Histogram showing the percentage frequency of Coulomb energy for 9 residues from-4 to +4 of HACs for Piwi, Su(var)3-9, Dmt, and XNP between 200 to 250 ns of the MD simulations, using the same format as C. (H) Coulomb energy ± SD [kJ/mol] of 15 HAC sequences from-7 to +7 between 200 to 250 ns of the MD simulations. The peptides are plotted according to ranking in terms of fold HP1a co-enrichment. The R-squared value is shown. (I) Histogram showing the percentage frequency of Coulomb energy from positions-7 to +7 of Piwi, Su(var)3-9, Dmt, and XNP HACs between 200 to 250 ns of the MD simulations, using the same format as C.

**Supplementary Figure 5:**
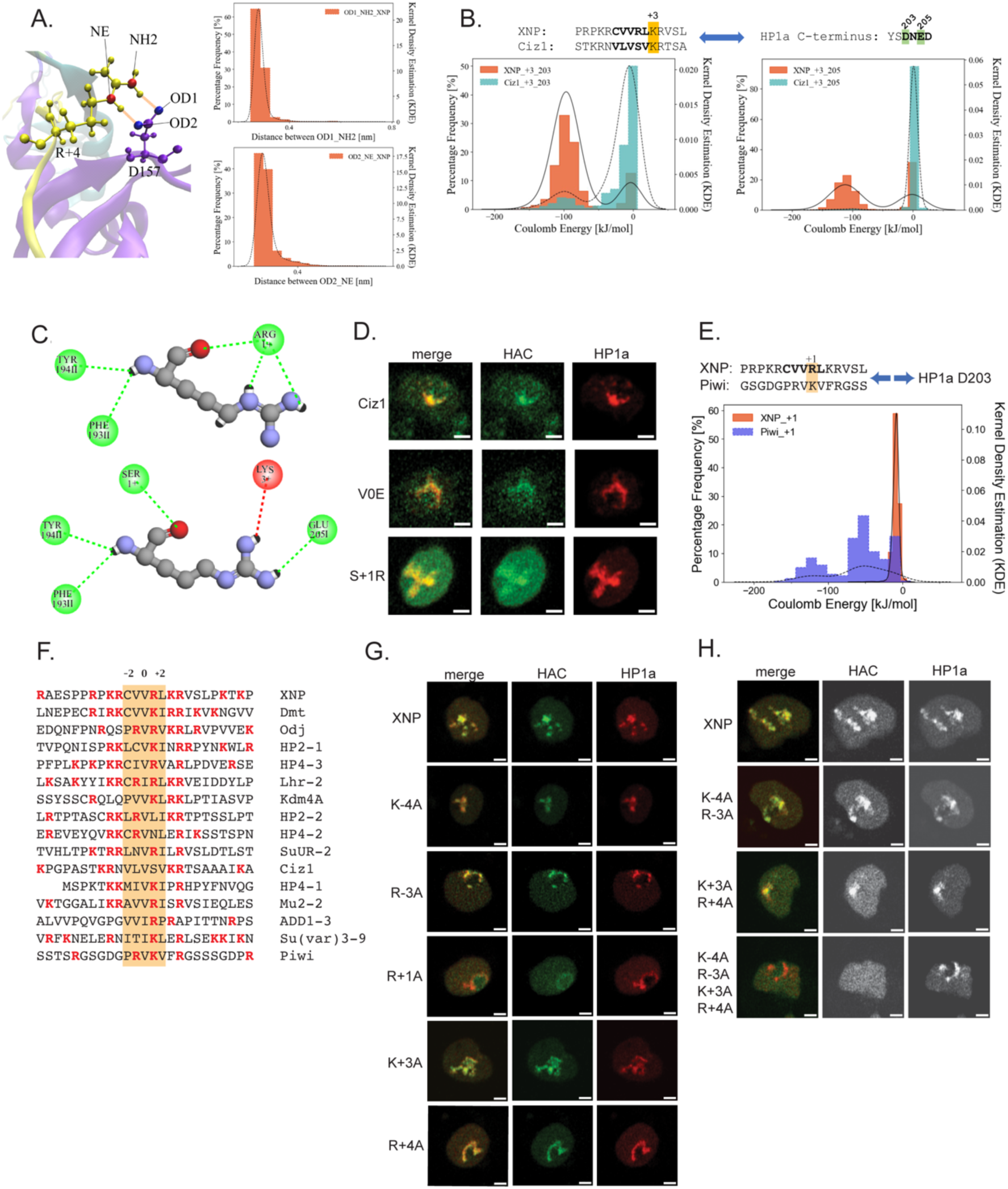
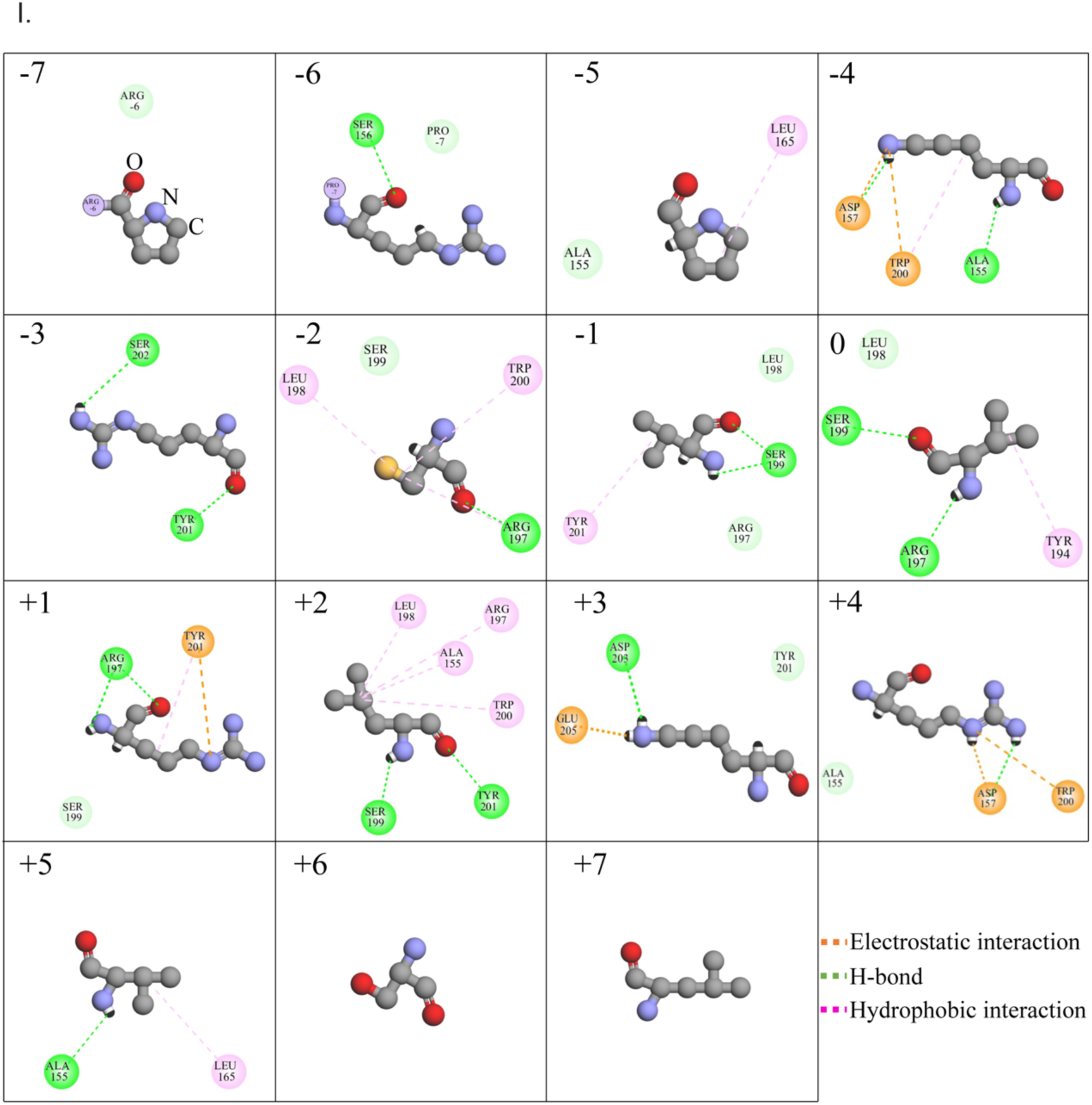
(A) Visualization of salt bridge interactions between XNP HAC Arg+4 and Glu157 of HP1a CSDII (left). Histograms of the percentage frequency of salt bridge formation between the guanidinium and a guanidino group of the XNP HAC Arg+4 sidechain and 2 O atoms of HP1a Glu157^II^ (right). (B) Histograms showing the percentage frequency of the salt bridges between Asp203^I^ and +3 (left) and between Glu205^I^ and +3 (right) of XNP as orange with a solid line, and of Ciz1 as cyan with a dashed line, between 200 to 250 ns of the MD simulations. (C) Interaction models of HP1a Arg197 showing 3 hydrogen bonds forming with Arg+1 of the XNP HAC (top), and a single hydrogen bond forming between Ser+1 plus charge repulsion with Lys+3 of the Ciz1 HAC (bottom). (D) Representative images of Ciz1 HAC and mutations either disabling central valine or adding a positive charge at the +1 position. (E) Histogram showing the percentage frequency of the Coulomb energy between Asp203^II^ or Ser199^I^ and +1 of XNP as orange with a solid line and Piwi as blue with a dashed line, between 200 to 250 ns of the MD simulations. All histograms were fitted by kernel density estimation (KDE). (F) Alignment of all 16 HACs with pentameric core highlighted in yellow and marked with-2, 0, and +2 positions, and positively charged residues marked in red. (G) Representative images of mNeonGreen-tagged XNP HACs with single alanine substitutions at positions-4,-3, +1, +3, or +4 co-transfected with mScarlet-I-tagged HP1a (H) Representative images of XNP HACs containing combinations of alanine mutations at either-4 and-3, +3 and +4, or all four positions. (I) Interactional profiles of XNP from-7 to +7 generated by Discovery Studio. Orange, green, and pink dashed lines represent an electrostatic interaction, hydrogen bond, and hydrophobic interaction. Grey, blue, and red atoms of amino acid structures indicate carbon, nitrogen, and oxygen, respectively.

**Supplementary Figure 6.**
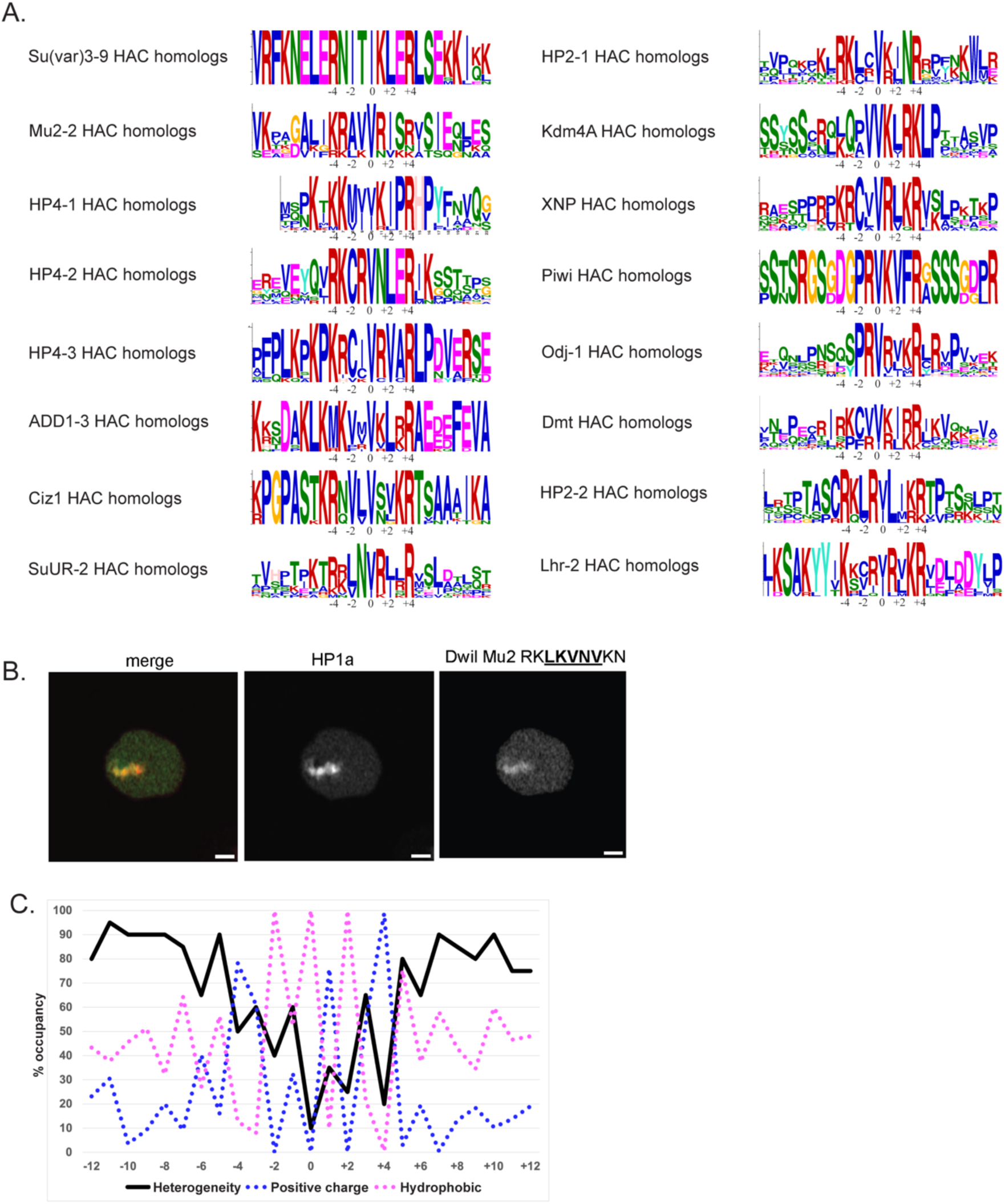
(A) HAC sequence pileup using MEME Suite 5.5.7 showing high conservation of specific residues among each group of homologs. (B) Imaging panels showing enrichment of putative Mu2 HAC from *Drosophila willistoni* containing a unique Asn+4. Scale bar = 1 um. (C) Positional bias for basic (red dots) and hydrophobic (blue dots) residues across Drosophila evolution of HACs is contrasted with the heterogeneity of amino acids (purple) for each position.

**Supplementary Figure 7.**
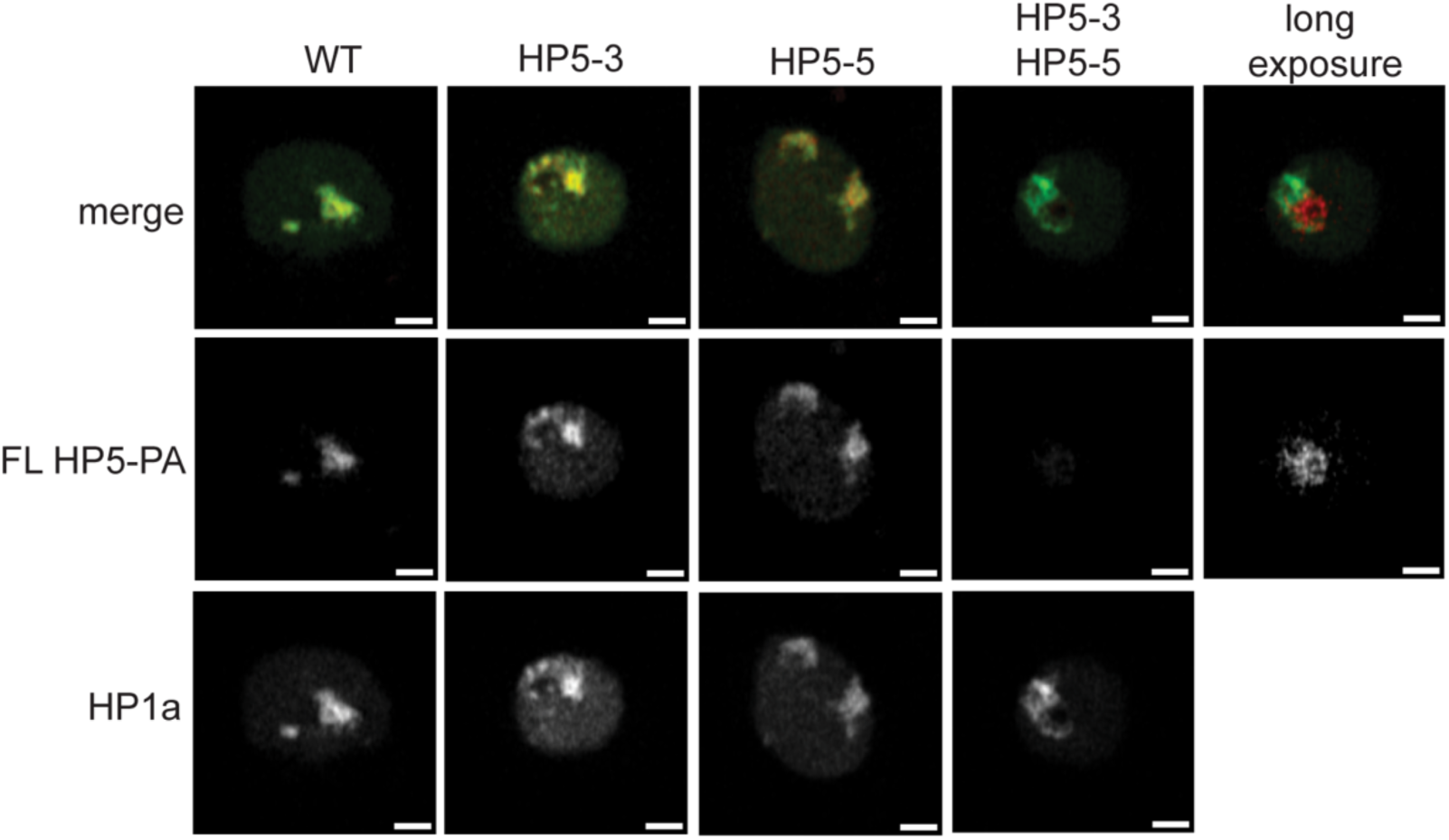
Representative images showing enrichment of wildtype or mutant full-length HP5-PA isoform at HP1a biocondensates. A longer exposure of the HP5 double mutant was necessary to visualize the loss of HP1a co-enrichment and accumulation at nucleoli. Scale bars = 2 microns.

**Supplementary Figure 8:**
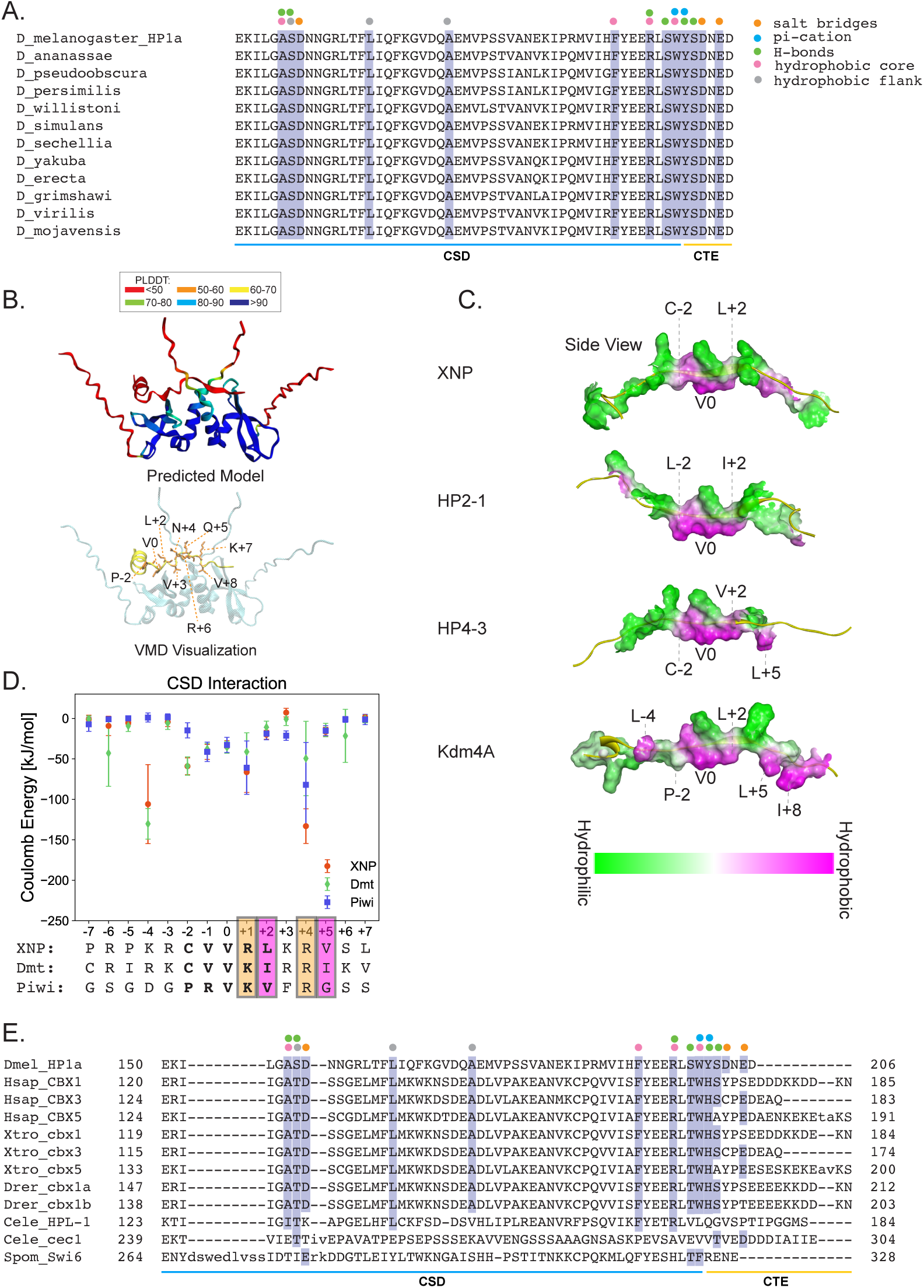
(A) Alignment of CSD and C-terminal tails of *D. melanogaster* HP1a homologs with other Drosophila species. Salt bridge-forming residues are marked with orange dots, pi-cation residues in blue, hydrogen bonds in green. Residues that interact with the hydrophobic core of HACs are marked with pink dots, whereas those that interact with flanking hydrophobic residues are marked with purple dots. Conservation is marked with purple highlight. HP1a CSD and CTE regions are marked with blue and yellow lines, respectively. (B) The Alphafold Multimer-predicted computational model (top) and VMD- visualized image of the model for the CSD dimer with C-terminal tails and ADD1-1 (bottom). The Alphafold Multimer-predicted structure is colored based on pLDDT (predicted local distance difference test) scores. Red: pLDDT<50, Orange: 50<pLDDT<60, Yellow: 60<pLDDT<70, Green: 70<pLDDT<80, Skyblue: 80<pLDDT<90, and Blue: 90<pLDDT.

**Supplementary Method Figure 1:**
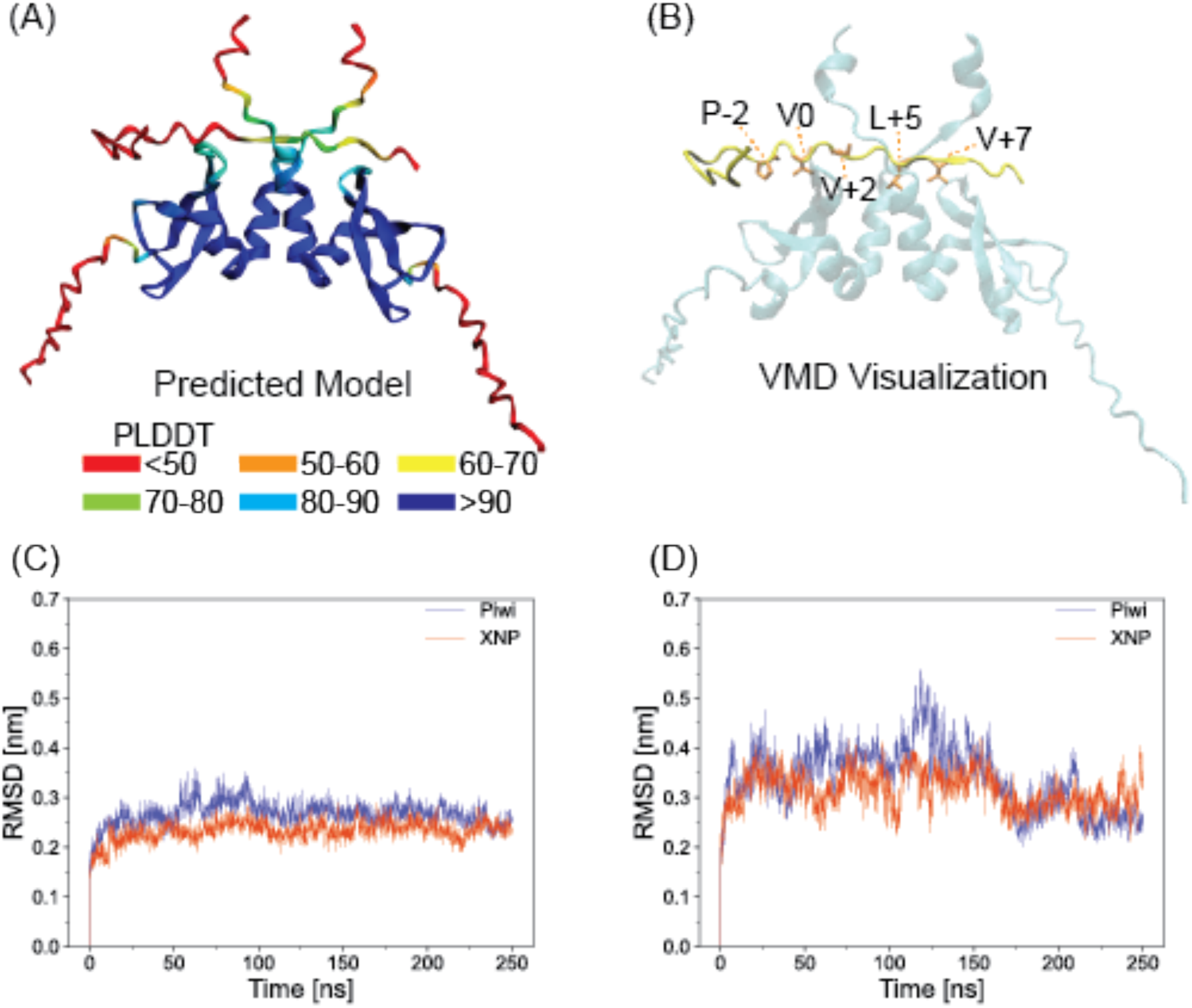
(A) The Alphafold Multimer-predicted computational model and (B) visualized image of the model for the CSD dimer with C-terminal tails and Odj-1. The Alphafold Multimer-predicted structure is colored based on pLDDT (predicted local distance difference test) scores. Red: pLDDT<50, Orange: 50<pLDDT<60, Yellow: 60<pLDDT<70, Green: 70<pLDDT<80, Skyblue: 80<pLDDT<90, and Blue: 90<pLDDT. CSD homodimer and the Odj-1 are colored cyan and yellow, respectively. Residues at the-2, 0, +2, +5, and +7 positions (in licorice drawing style) are colored orange, with the amino acid code indicated. (C and D) Root mean square deviation (RMSD) of (C) the HP1a CSD dimer (amino acids 141-201) in Piwi and XNP peptide models and (D) the HAC peptides of Piwi and XNP from-7 to +7 amino acids throughout the 250 ns simulations.

**Supplementary Method Table 1:** The 25-amino-acid sequences of 16 heterochromatin-enriched peptides, with the central valine (V) designated as position 0. The table also includes AlphaFold-Multimer prediction scores: pLDDT (predicted Local Distance Difference Test, scaled from 0 to 100) and pTM (predicted Template Modelling, scaled from 0 to 1) were shown.

| Peptide Name | Amino Acid Sequences |  |  |  |  |  |  |  |  |  |  |  |  |  |  |  |  |  |  |  |  |  |  |  |  | Prediction Scores |  |
| --- | --- | --- | --- | --- | --- | --- | --- | --- | --- | --- | --- | --- | --- | --- | --- | --- | --- | --- | --- | --- | --- | --- | --- | --- | --- | --- | --- |
|  | -12 | -11 | -10 | -9 | -8 | -7 | -6 | -5 | -4 | -3 | -2 | -1 | 0 | 1 | 2 | 3 | 4 | 5 | 6 | 7 | 8 | 9 | 10 | 11 | 12 | pLDDT | pTM |
| XNP | R | A | E | S | P | P | R | P | K | R | C | V | V | R | L | K | R | V | S | L | P | K | T | K | P | 76.7 | 0.706 |
| Dmt | L | N | E | P | E | C | R | I | R | K | C | V | V | K | I | R | R | I | K | V | K | N | G | V | V | 75.9 | 0.696 |
| Odj-1 | E | D | Q | N | F | P | N | R | Q | S | P | R | V | R | V | K | R | L | R | V | P | V | V | E | K | 72 | 0.655 |
| HP2-1 | T | V | P | Q | N | I | S | P | R | K | L | C | V | K | I | N | R | R | P | Y | N | K | W | L | R | 77.3 | 0.729 |
| HP4-3 | P | F | P | L | K | P | K | P | K | R | C | I | V | R | V | A | R | L | P | D | V | E | R | S | E | 76.2 | 0.705 |
| Lhr-2 | L | K | S | A | K | Y | Y | I | K | R | C | R | I | R | L | K | R | V | E | I | D | D | Y | L | P | 80.1 | 0.761 |
| Kdm4A | S | S | Y | S | S | C | R | Q | L | Q | P | V | V | K | L | R | K | L | P | T | I | A | S | V | P | 75.7 | 0.714 |
| HP2-2 | L | R | T | P | T | A | S | C | R | K | L | R | V | L | I | K | R | T | P | T | S | S | L | S | L | 77.1 | 0.732 |
| HP4-2 | E | R | E | V | E | Y | Q | V | R | K | C | R | V | N | L | E | R | I | K | S | S | T | S | P | N | 75.6 | 0.687 |
| SuUR-2 | T | V | H | L | T | P | K | T | R | R | L | N | V | R | I | L | R | V | S | L | D | T | L | S | T | 77.6 | 0.735 |
| Ciz1 | K | P | G | P | A | S | T | K | R | N | V | L | V | S | V | K | R | T | S | A | A | A | I | K | A | 75.4 | 0.719 |
| HP4-1 |  |  |  | M | S | P | K | T | K | K | M | I | V | K | I | P | R | H | P | Y | F | N | V | Q | G | 78.6 | 0.727 |
| Mu2-2 | V | K | T | G | G | A | L | I | K | R | A | V | V | R | I | S | R | V | S | I | E | Q | L | E | S | 77.8 | 0.732 |
| ADD1-3 | K | K | S | D | A | K | L | K | M | K | V | M | V | K | L | K | R | A | E | D | E | F | E | V | A | 79.4 | 0.741 |
| Su(var)3-9 | V | R | F | K | N | E | L | E | R | N | I | T | I | K | L | E | R | L | S | E | K | K | I | K | N | 78.2 | 0.74 |
| Piwi | S | S | T | S | R | G | S | G | D | G | P | R | V | K | V | F | R | G | S | S | S | G | D | P | R | 75.4 | 0.688 |

## Notes

### Competing Interest Statement

The authors have declared no competing interest.

### Summary of Updates

Authors were revised, plus minor details in the text.

